# α-Synuclein pathology in Parkinson disease activates homeostatic NRF2 anti-oxidant response

**DOI:** 10.1101/2021.04.15.439988

**Authors:** Alberto Delaidelli, Mette Richner, Lixiang Jiang, Amelia van der Laan, Ida Bergholdt Jul Christiansen, Nelson Ferreira, Jens R. Nyengaard, Christian B. Vægter, Poul H. Jensen, Ian R. Mackenzie, Poul H. Sorensen, Asad Jan

## Abstract

Circumstantial evidence points to a pathological role of alpha-synuclein (aSyn; gene symbol *SNCA*), conferred by aSyn misfolding and aggregation, in Parkinson disease (PD) and related synucleionpathies. Several findings in experimental models implicate perturbations in the tissue homeostatic mechanisms triggered by pathological aSyn accumulation, including impaired redox homeostasis, as significant contributors in the pathogenesis of PD. The nuclear factor erythroid 2-related factor (NRF2) is recognized as ‘the master regulator of cellular anti-oxidant response’, both under physiological as well as in pathological conditions. Using immunohistochemical analyses, we show a robust nuclear NRF2 accumulation in post-mortem PD midbrain, detected by NRF2 phosphorylation on serine residue 40 (nuclear active p-NRF2, S40). Curated gene expression analyses of four independent publicly available microarray datasets revealed considerable alterations in NRF2-responsive genes in the disease affected regions in PD, including substantia nigra, dorsal motor nucleus of vagus, locus coeruleus and globus pallidus. To further examine the putative role of pathological aSyn accumulation on nuclear NRF2 response, we employed a transgenic mouse model of synucleionopathy (M83 line, Prnp-SNCA*Ala53Thr), which manifest widespread aSyn pathology (phosphorylated aSyn; S129) in the nervous system following intramuscular inoculation of exogenous fibrillar aSyn. We observed strong immunodetection of nuclear NRF2 in neuronal populations harboring p-aSyn (S129), and found an aberrant anti-oxidant and inflammatory gene response in the affected neuraxis. Taken together, our data support the notion that pathological aSyn accumulation impairs the redox homeostasis in nervous system, and boosting neuronal anti-oxidant response is potentially a promising approach to mitigate neurodegeneration in PD and related diseases.

## INTRODUCTION

Parkinson disease (PD) is a major neurodegenerative cause of chronic dysfunction in the subcortical somatomotor system, which is frequently compounded by non-motor symptoms of extranigral origin(1–3). The disease is clinically characterized by the cardinal features of resting tremor, bradykinesia, rigidity and gait apraxia(1, 2). The neuropathology of PD is principally defined by the loss of dopaminergic neurons in the midbrain *substantia nigra* (SN)-*pars compacta*, and immunoreactive deposits of α-synuclein (aSyn; gene symbol *SNCA*) protein across multiple regions in the central nervous system (CNS)(1, 2, 4). Pathological aSyn deposition in the CNS, termed Lewy body (LB) pathology, is also seen in other neurodegenerative diseases such as Alzheimer disease (AD), Dementia with Lewy bodies (DLB), and in Multiple system atrophy (MSA)(5). In the vast majority of PD cases, the disease is of idiopathic (non-inheritable) origin, and genetic factors underlie 5-10% of clinically diagnosed PD(2). Rare missense mutations in *SNCA* that result in N-terminal amino acid substitutions in aSyn, or multiplications in *SNCA* gene locus are recognized etiological factors in the autosomal-dominant forms of PD(2, 6, 7). Additionally, mutations in several other genes (of autosomal-dominant or recessive inheritance) have been discovered to cause rare forms of familial PD, underscoring the complex etiology of the disease(1, 2). Furthermore, the occurrence of distinct neurodegenerative lesions and progressive aSyn pathology in PD point to the selective vulnerability of specific subcortical nuclei, with relative sparing of other brain regions(4, 8, 9).

Experimental models based on the genetic aberrations in familial PD have revealed a number of candidate mechanisms that are potentially relevant to neurodegeneration, and for the development of mechanism-based therapies in PD. These studies implicate perturbed cellular homeostasis caused by several biochemical alterations including defective autophagic flux, endoplasmic reticulum (ER) stress, calcium dyshomeostasis and mitochondrial impairment(10–13). Although predominantly studied as a sequelae to aSyn aggregation (10, 11), the relevance of these mechanisms is also supported by findings in experimental paradigms based on other genetic factors causing PD. For instance, mutations in leucine-rich repeat kinase 2 (*LRRK2*)-the most common cause of the autosomal dominant PD- are associated functional alterations such as impaired vesicular trafficking and cytoskeleton dynamics, defective autophagy and lysosomal degradation, and mitochondrial dysfunction(14, 15). Furthermore, mutations of Parkin (*PARK2*; an E3 ubiquitin ligase), the protein deglycase DJ-1 (*PARK7*) and the PTEN-induced putative kinase 1 (*PINK1*) cause functional deficits affecting the autophagic flux of damaged mitochondria and neuroprotective response against oxidative stress(12). Moreover, inadequate mitochondrial complex I activity, as a result of aSyn aggregation or in chemically induced experimental parkinsonism, has been implicated in the impaired energy production, and in the oxidative damage due to increased free radical production(16, 17). Similarly, post-mortem studies in PD implicate aberrant ROS homeostasis, lipid peroxidation, protein nitration and nucleic acid oxidation as significant disease associations(18, 19). Lastly, cellular oxidative stress has been shown to exacerbate aSyn misfolding and aggregation, thus triggering a vicious circle that culminates in cytotoxicity and further redox dyshomeostasis(20). Although, the nature of the final common pathway(s) that triggers neurodegeneration in PD remains debatable, an interplay of the above mentioned genetic factors and environmental triggers are thought to blunt the protective tissue homeostatic response against the deleterious effects of proteotoxic and oxidative stress(11, 12).

In this context, the nuclear factor erythroid 2–related factor 2 (protein, NRF2/Nrf2; gene symbol *NFE2L2*), a member of basic leucine zipper (bZIP) protein family, plays a crucial role in the cellular adaptive response under oxidative and metabolic stress(21, 22). Under resting metabolic conditions, NRF2 is sequestered in the cytoplasm by its inhibitor, Kelch-Like ECH-Associated Protein-1 (protein, Keap1; gene symbol *KEAP1*), which also mediates the proteosomal degradation of NRF2(22). A mismatch between the production of reactive oxygen species (ROS) free radicals and ROS scavenging mechanisms induces the phosphorylation of NRF2 at serine 40 residue (S40) and disrupts the NRF2-Keap1 complex. As a result, there is increased cytoplasm-to-nucleus translocation of NRF2, and subsequent transactivation of the pathways involved in antioxidant, anti-inflammatory and xenobiotic defence (including NRF2 auto-regulation)(22). In addition to the canonical regulation via Keap1 mediated autophagic degradation, NRF2 activity is also influenced by several factors including post-translational modifications, epigenetic factors, microRNAs and metabolic adaptations under nutrient stress(22–24). Burgeoning evidence points to the relevance of aberrant NRF2 activity in the pathogenesis of neurodegenerative diseases, including PD(25, 26). First, there is a correlative decline in NRF2 activity with age, which is the strongest risk factor in the common neurodegenerative diseases(27). Second, neuropathological studies suggest increased nuclear localization of total NRF2 in the post-mortem substantia nigra of PD patients, in contrast to the predominant cytosolic localization in controls(25). Third, single nucleotide polymorphisms (SNPs) in *NFE2L2* or the promoter region are associated with altered disease risk or the age of PD onset(28). Therefore, understanding the role of NRF2 response in the pathophysiology of neurodegeneration in PD and related diseases, and investigating the therapeutic impact of boosting NRF2 anti-oxidant response has gained significant interest(16, 26, 28).

In this report, we show increased nuclear localization of NRF2 in post-mortem PD midbrain in the presence of Lewy related aSyn pathology, as detected by immunonhistochemical (IHC) detection of phosphorylated NRF2 on residue serine-40 (p-NRF2, S40)- a robust posttranslational modification associated with the nuclear accumulation of NRF2(29–31). By analyzing four independent microarray studies in PD (32–35), we also report altered expression of NRF2-responsive genes in PD affected brain regions. In order to establish the role of pathological aSyn accumulation to altered NRF2 response, we performed IHC and gene expression analyses in a transgenic mouse model of synucleinopathy expressing the PD-associated mutant *Ala53Thr* (A53T) aSyn (the M83 line)(36, 37). In this rodent model, widespread aSyn pathology (phosphorylated aSyn, serine- 129; p-aSyn, S129) in the CNS is ectopically induced by the intramuscular inoculation of pre-formed fibrillar (PFF) aSyn in the hindlimb. Specifically, the intracerebral aSyn pathology is predominantly observed in the brainstem regions around 50-70 days post-injection, with relative paucity in the forebrain areas(36, 37). Our data show considerable enrichment of nuclear NRF2 (p- NRF2, S40) in the brain regions harboring aSyn pathology (p-aSyn, S129), and distinct alterations in the NRF2-dependent anti-oxidant and inflammatory gene expression response in this prion-like model of synucleionoapthy. Taken together, our data provide fruitful insights into the putative role of NRF2 stress adaptive signaling in PD in relation to pathological aSyn accumulation, and strengthen the rationale for boosting NRF2-anti-oxidant response towards mitigating the deleterious effects of proteopathic stress.

## RESULTS

### IHC analyses show increased nuclear localization of phosphorylated NRF2 (p-NRF2, S40) in post-mortem PD midbrain

Previously, it has been reported that in PD susbtantia nigra, NRF2 is abundantly detected in both cytosolic and nuclear locations, and exhibited relatively higher localization in the neuronal nuclei than was observed in control brains(25). However, it remains to be determined if the nuclear enrichment of NRF2 in PD also correlates with its cytoprotective and anti-oxidant response. NRF2 is highly expressed in neuronal and glial cells(38), and its stability and nuclear transcription activity are regulated by several factors including post-translational modifications such as phosphorylation and acetylation(31). Among the phosphoryaltion sites, serine-40 (p-S40) has been used in several studies as a surrogate marker of NRF2 stability and its nuclear localization, and remains the most widely studied post translational modification in the published literature to date(31, 39). Although the identification of putative kinase(s) represents an evolving field, experiments involving acute exposure of cultured cells to mitochondrial ROS inducers and site-directed mutagenesis approaches implicate a role of protein kinase C (PKC)(29, 40).

Accordingly, we performed IHC analyses of phosphorylated NRF2 (p-NRF2, S40) in postmortem midbrain sections obtained from controls and PD cases (Table S1). In parallel, using serial sections, we also assessed the phosphorylation of aSyn on serine residue 129 (p-aSyn, S129), which is a widely used neuropathological marker for detecting Lewy related aSyn pathology in tissue specimen(5, 36, 41, 42). Our IHC data revealed distinct nuclear p-NRF2 (S40) in PD brains compared with control brains, both in substantia nigra (SN) and periaqueductal grey (PAG) regions (Compare Fig. 1A-B, Controls and 1C-D, PD; also see Fig. 1E-F and S1A-B). aSyn LB pathology (detected by p-aSyn, S129 IHC) was also conspicuous in SN and PAG (Fig. 1C, 1E and S1B), and is a characteristic feature of PD(2, 4, 41).

**Figure 1.**
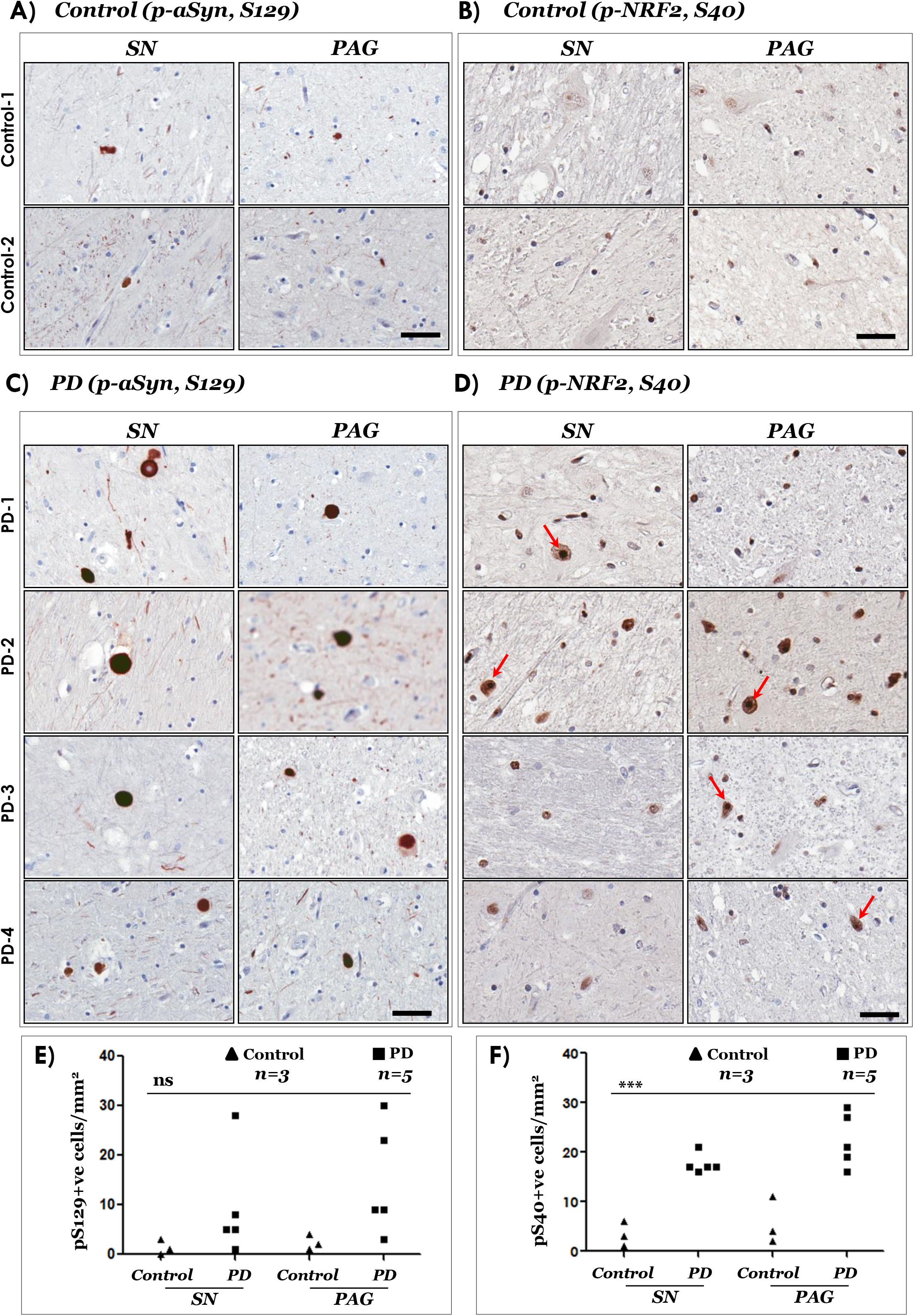
Immunostaining of phospho-aSyn (S129) and phospho-NRF2 (S40) in postmortem control and PD midbrain sections. **(A-D)** Representative IHC images showing phospho-aSyn (S129) and phospho-NRF2 (S40) immunostaining in substantia nigra (SN) and periaqueductal grey (PAG) of two controls (A-B) and four PD cases (C-D). Red arrows in the 20X magnified views point to cells with predominant nuclear localization of phospho-NRF2 (scale bar=100 μm). Also see Figure S1 showing panoramic and magnified views from a control and a PD case and Table S1. Primary antibodies: p-aSyn (S129)- 81A (in A and C) and p-NRF2 (S40)- EP1809Y (in B and D). **(E-F)** Semi-quantitative analyses of p-aSyn (pS129; in E) and p-NRF2 (pS40; in F) immunopositive cells in the indicated regions of control and PD midbrain sections. Individual data points represent immunopositive cell counts/mm^2^/region in each section, with controls being depicted as black triangles and PD as the black squares. (SN, substantia nigra; PAG, periaqueductal grey matter; One-way ANOVA: ns, not significant; ***p<0.0001).

### Microarray gene expression analyses indicate aberrant NRF2 anti-oxidant response and activation of pro-apoptotic factors in PD

Next, we investigated the functional significance of the observations concerning the increased NRF2 nuclear localization (p-NRF2, S40; Fig. 1 and S1) in PD brain. In particular, we focused on assessing the expression of NRF2-responsive ROS detoxification factors (including auto-regulation), and the expression of pro-apoptotic caspases implicated in neuronal loss in PD(43). For this purpose, we queried four publicly available functional genomics datasets available in the GEO repository of NCBI(32–35, 44). Following is the summary of data with noticeable differences between control and PD tissues: First, *NFE2L2* (encoding NRF2) showed increased expression in in PD SN and locus coeruleus-LC with a higher (albeit, statistically not significant) expression in the dorsal motor nucleus of vagus- DMX (Fig. 2A, GSE4349). Intriguingly, the expression of *KEAP1* (NRF2 inhibitor) was also found to be highly increased in PD SN (Fig. 2B, GSE7621; S3B, GSE26297). Second, among the NRF2 target gens involved in the anti-oxidant pathways, the expression of heme oxyganase 1 (gene symbol, *HMOX1*; protein, HO-1; involved in heme catabolism and ROS detoxification) was increased in PD SN and globus pallidus interna- GPi (Fig. 2C, GSE7621; GSE20146). Similarly, the expression of gamma-glutamylcysteine synthetase (gene symbol, *GCLC*; a rate-limiting enzyme in the anti-oxidant glutathione synthesis) was also induced in PD SN (Fig. 2D, GSE43490). Lastly, we also probed if the local changes in NRF2 homeostatic response are potentially associated with factors regulating apoptosis, in particular caspases. Accordingly, we found higher expression of the pro-apototic caspase-3 (*CASP3*) in all the datasets queried; albeit, the differences were statistically significant only in GSE43490 (Fig. 2E, SN, DMX and LC). Similarly, the executioner pro-apoptotic caspase 6 (*CASP6*) was also found to be significantly altered in PD SN (Fig. 2F, GSE43490).

**Figure 2.**
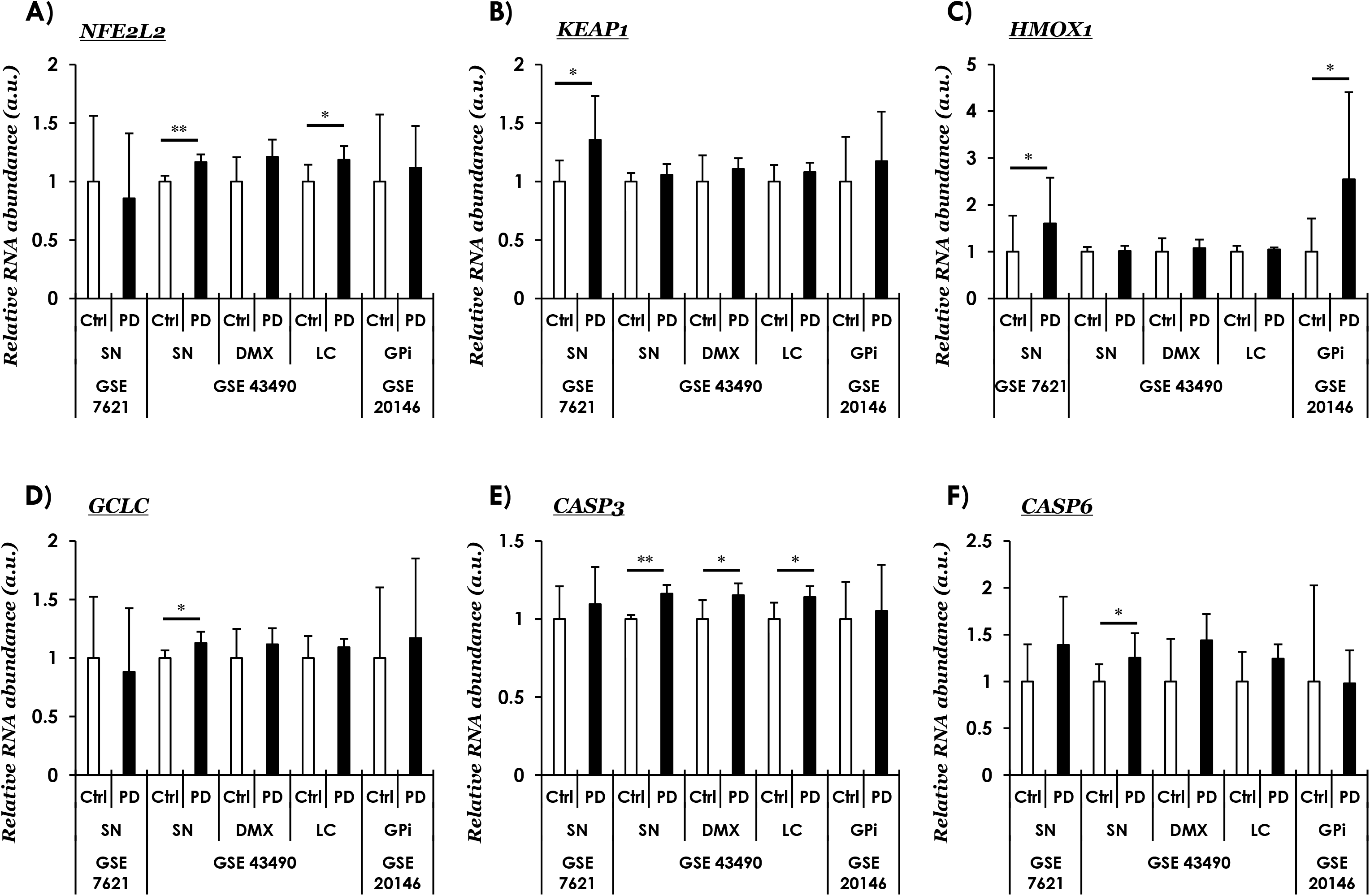
Curated gene expression analyses of publicly available microarray datasets in PD using Gene Expression Omnibus (GEO). **(A)** *NFE2L2* (Nuclear Factor, Erythroid 2 Like 2, NRF2)**, (B)** *KEAP1* (Kelch Like ECH Associated Protein 1, NRF2 inhibitor protein), **(C-D)** NRF2 anti-oxidant response mediators, *HMOX1* (Heme Oxygenase 1, in C), *GCLC* (Glutamate-Cysteine Ligase Catalytic Subunit, in D), and **(E-F)** Pro-apoptotic caspases, *CASP3* (Caspase 3, in E) and *CASP6* (Caspase 6, in F). The values across the datasets are expressed relative to the controls in each microarray dataset, i.e., mean value of control samples=1 (a.u., arbitrary units). Error bars represent standard deviation of the mean, s.d. In GSE7621- SN (*substantia nigra*); controls (Ctrl, n=9) and PD (Parkinson Disease cases, n=16); in GSE43490- SN (*substantia nigra*; controls, n=6 and PD, n=8), DMX (*dorsal motor nucleus of vagus*; controls, n=5 and PD, n=8), LC (*locus coeruleus*; controls, n=7 and PD, n=8); in GSE20146- GPi (*globus pallidus interna*; controls, n=10 and PD, n=10). Pair-wise comparisons were assessed by Mann-Whitney test- only significant differences (*=p≤0.05, **=p≤0.01) are highlighted. Probe IDs, microarray platforms and source studies are listed in Table S2. Also see Figure S2-S3.

We have also probed the expression of additional NRF2-responsive genes, including NAD(P)H Quinone Dehydrogenase 1 (*NQO1*- involved in xenobiotic detoxification), neuroprotector molecules such as the putative NRF2 stabilizer protein deglycase DJ1 (*PARK7*, mutated in subset of familial PD) and inflammatory mediators interleukin-1 (*IL1*) and the tumor necrosis factor (*TNF*) (S2). Among these, only *NQO1* showed a significant change in PD GPi (S2C, GSE20146). The dataset GSE26927(34), shown in S3, also contains expression profiles of additional neurodegenerative diseases beside PD, in which defective ROS metabolism is implicated. Keeping in view a broad interest in the topic, we have summarized the data included in GSE26927 for readers (S3); however, their elaboration is beyond the scope of this report. Collectively, these data (Fig. 2 and S2) support the notion regarding altered redox homeostasis in PD. Nevertheless, it is also noteworthy that the expression of a given gene does not show significant differences in all the examined datasets in PD (elaborated in Discussion).

### Brainstem α-synucleinopathy in M83 mice leads to increased phospho-NRF2 (S40) nuclear accumulation

In order to assess the significance of pathological CNS aSyn accumulation in the context of the altered NRF2-dependent gene response observed in PD, we examined phospho-NRF2 (S40) immunostaining, in conjunction with the assessment of NRF2-responsive gene expression in brains of transgenic M83 mice (expressing the aggregation prone A53T mutant human aSyn)(37). Bilateral intramuscular inoculation of mouse PFF aSyn into the hindlimb of M83 mice results in severe morbidity and senosorimotor defects, initially characterised by a unilateral footdrop which progresses to complete paralysis 8-10 weeks postinjection(36). The genesis of CNS aSyn neuropathology and sensorimotor deficits in this PFF based model of peripheral-to-central propagation of syncleinopathy are extensively characterized by several laboratories including our own(36, 45–47). For these studies, we used tissues collected from terminal stage homozygous M83^+/+^ mice, once a unilateral foot drop was clearly established (70-90 days post-PFF aSyn inoculation)(36, 46).

In line with the previous reports, we also found predominant aSyn pathological affection of brainstem nuclei involved in locomotor control, and also in the periaqueductal grey (PAG) (36, 47). Accordingly, using immunostaining for p-aSyn (S129), we found abundant aSyn pathology in PAG, red nucleus, pontine gigantocellualr nuclei (Gi) and pontine vestibular nuclei (VN) (Fig. 3A and S6A). Among the additional regions examined, aSyn (p-aSyn, S129) accumulation was relatively sparse (Fig. 3A, midbrain tegmentum- containing substantia nigra- and frontal cortex), or largely undetectable (Fig. 3A, cerebellar lobules cb1-5 and striatum), as has also been shown(36, 47). Additionally, p-aSyn (S129) accumulation in the brains from PBS (vehicle) injected cohort was not remarkable (S4A). This is not surprising as naive M83 mice (i.e, not injected with PFF aSyn) develop spontaneous aSyn pathology usually after 8-12 months of age, and it is extremely rare before 7 months of age(36, 37).

**Figure 3.**
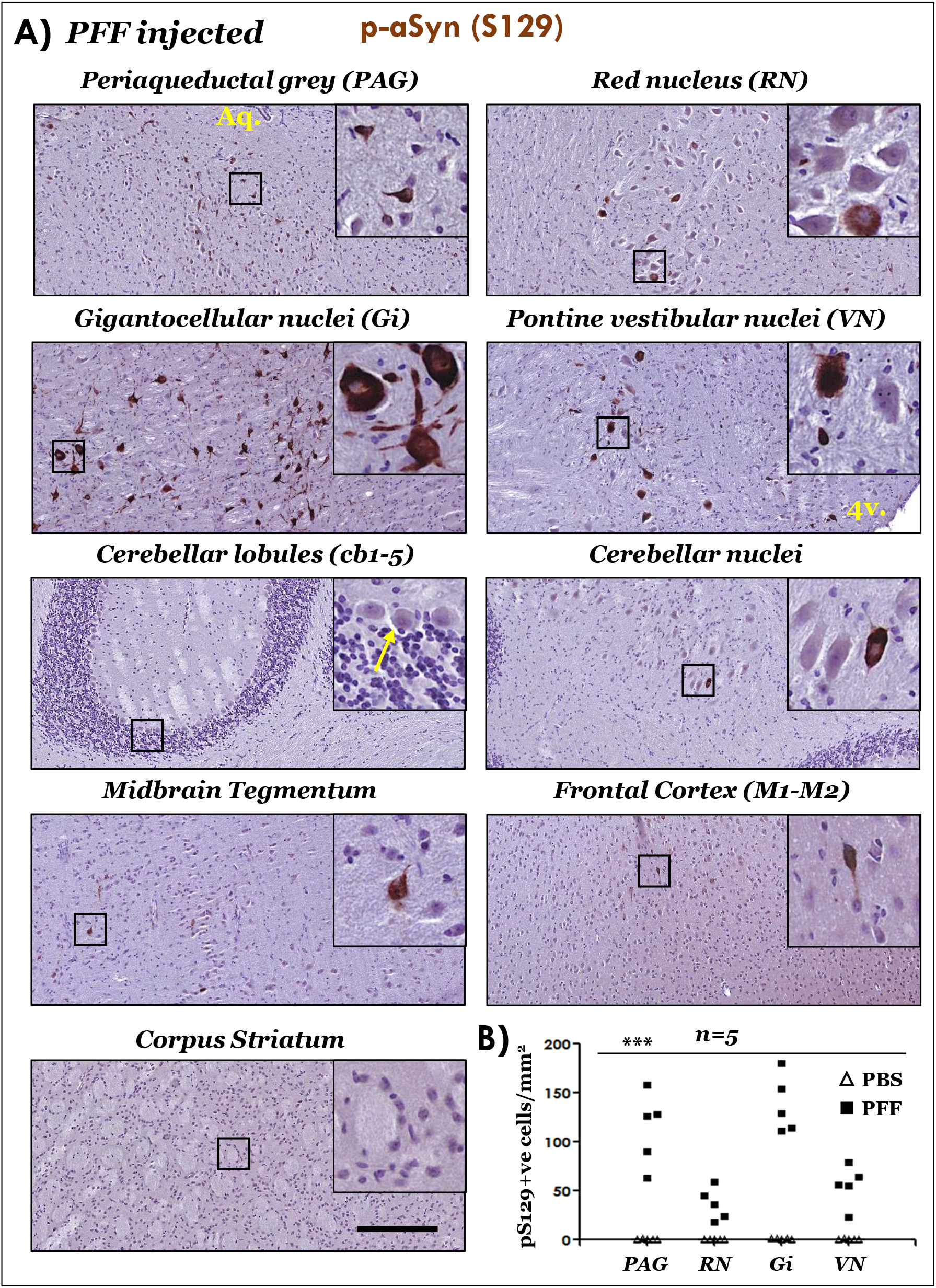
Immunostaining of phospho-aSyn (S129) in the brain regions of PFF aSyn injected M83^+/+^ mice. **(A)** Representative IHC images showing phospho-aSyn (S129) immunostaining in neuronal somata and processes in the indicated brain regions. Also, notice the scarcity of staining in cerebellar lobules cb1-5 (purkinje and granule cells- yellow arrow in the inset), motor cortex and corpus striatum (10X low magnification views and 40X magnified views in the insets; scale bar=200 μm; Aq. in the image showing PAG, cerebral aqueduct; 4v. in the image showing vestibular nuclei image, 4^th^ ventricle). Bregma co-ordinates for the brain regions were determined, according to Paxinos and Franklin: (Bregma, −3.87 mm) midbrain at the level of superior colliculi showing PAG, red nucleus and tegmentum; (Bregma, −5.99) pontocerebellar junction showing cerebellar nuclei, cerebellar lobules (*cb1-5*), vestibular nuclei and pontine gigantocellualr nuclei (Gi); and (Bregma, 0.49 mm) forebrain showing motor cortex (M1 and M2) and corpus striatum). Also see Figure S4A (PBS injected cohort) and Figure S6A (additional high resolution data from the PFF injected cohort). Primary antibody in A: p-aSyn (S129)- 11A5. **(B)** Semi-quantitative analyses of p-aSyn (pS129) immunopositive cells in the indicated brain of PBS or PFF injected M83^+/+^ mice. Individual data points represent p-aSyn (S129) cell counts/mm^2^/region in each animal of the respective cohort, with PBS cohort being depicted as blank triangles and PFF as the black squares. (PAG, periaqueductal grey matter; RN, red nucleus; Gi, pontine gigantocellualr nuclei; VN, pontine vestibular nuclei; One-way ANOVA, ***p<0.0001; n=5/group).

Next, we assessed the nuclear NRF2 accumulation, by the IHC immunodetection of p-NRF2 (S40) in brain sections, from PBS and PFF aSyn injected cohorts. Robust nuclear p-NRF2 (S40) accumulation was observed in PAG, RN, Gi and VN, regions with significant p-aSyn (S129) accumulation in the PFF injected mice (Fig. 4A, also see S5A and S6B-C). Furthermore, in the regions with sparse aSyn pathology, e.g., midbrain tegmentum and frontal cortex, clear distinction (i.e., nuclear vs cytosolic) of p-NRF2 (S40) immunostaining was less pronounced, compared to the aSyn pathology affected regions (Fig. 4A). Lastly, in the cerebellar nuclei (with sparse aSyn pathology), nuclear p-NRF2 (S40) immunostaining was completely lacking and/or not consistent (Fig. 4A; also see S6B-C). This underlines potential differences in baseline and/or inducible degree of NRF2 response in distinct neuronal populations, and could- among other factors- represent an inherent property of their reserve for stress adaption (see Discussion).

**Figure 4.**
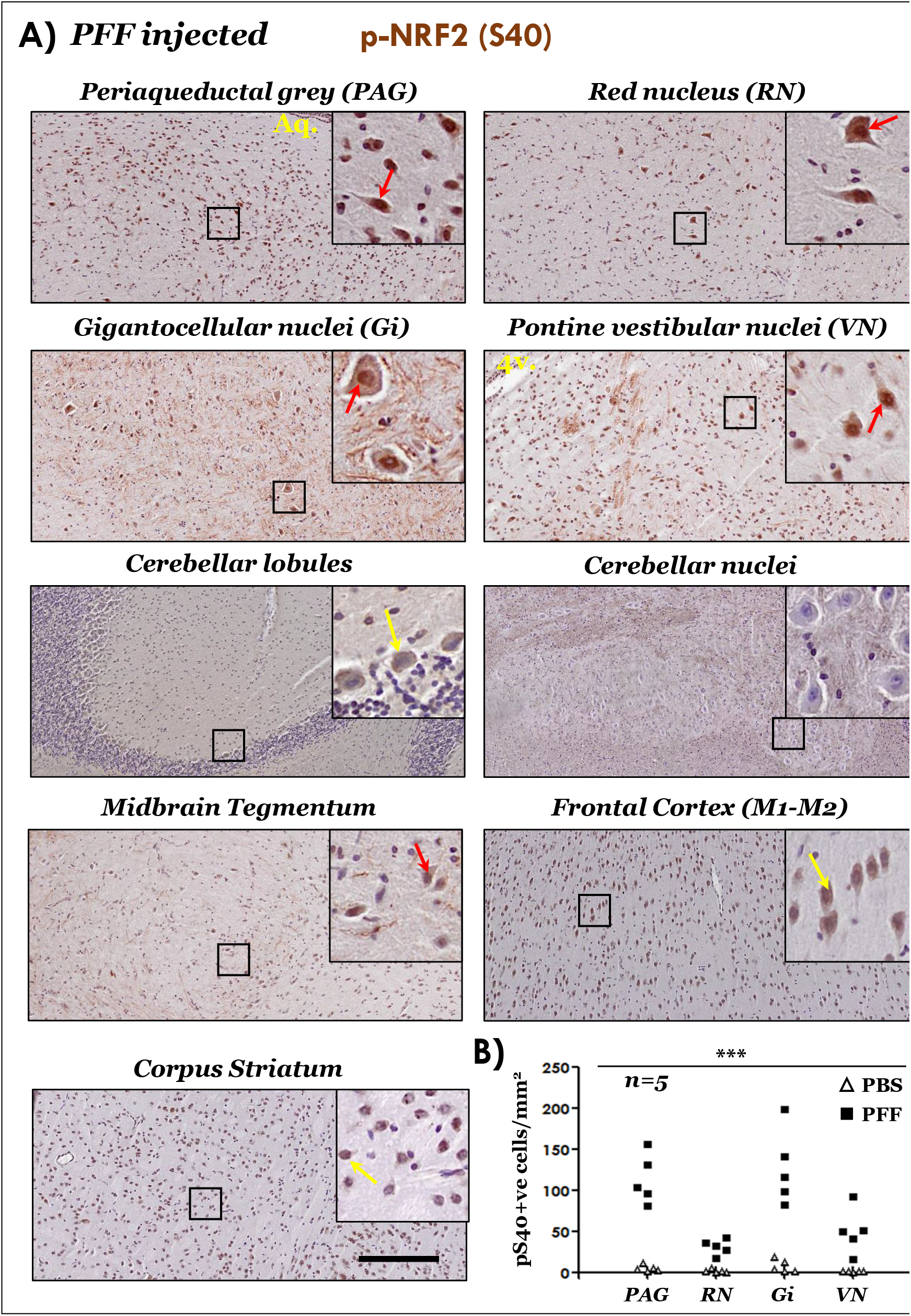
Immunostaining of phospho-NRF2 (S40) in the brain regions of PFF aSyn injected M83^+/+^ mice. **A)** Representative IHC images showing phospho-NRF2 (S40) immunostaining with distinct nuclear staining in brainstem regions (red arrows). Also, notice the predominantly cytoplasmic staining in cerebellar lobules (purkinje cells), motor cortex and corpus striatum- yellow arrows (10X low magnification views and 40X magnified views in the insets; scale bar=200 μm; Aq. in the image showing PAG, cerebral aqueduct; 4v. in the image showing vestibular nuclei image, 4^th^ ventricle). Bregma co-ordinates for the brain regions were determined, according to Paxinos and Franklin: (Bregma, −3.63 mm) midbrain at the level of superior colliculi showing PAG, red nucleus and tegmentum; (Bregma, −5.79) pontocerebellar junction showing cerebellar nuclei, cerebellar lobules (*cb1-5*), vestibular nuclei and pontine gigantocellualr nuclei (Gi); and (Bregma, 0.73 mm) forebrain showing motor cortex (M1 and M2) and corpus striatum. Also see Figure S5A (PBS injected cohort) and Figure S6B-C (additional high resolution data from the PFF and PBS cohorts). Primary antibody in A: p-NRF2 (S40)- PA5-67520. **(B)** Semi-quantitative analyses of p-NRF2 (pS40) immunopositive cells in the indicated brain regions of PBS or PFF injected M83^+/+^ mice. Individual data points represent p-NRF2 (pS40) cell counts/mm^2^/region in each animal of the respective cohort, with PBS cohort being depicted as blank triangles and PFF as the black squares. (PAG, periaqueductal grey matter; RN, red nucleus; Gi, pontine gigantocellualr nuclei; VN, pontine vestibular nuclei; One-way ANOVA, ***p<0.0001; n=5/group).

### α-syn pathology is associated with altered cytoprotective gene response in the brains of M83 mice

Next we wanted to determine if the changes in nuclear localization of NRF2 (p-NRF2, S40) in association with pathological aSyn accumulation leads to NRF2-dependent tissue response, i.e., the expression of anti-oxidant and cytoprotective genes. For this purpose, we assessed gene expression (by qRT-PCR) across the brain neuraxis, both in regions harboring significant aSyn pathology (i.e., brainstem), as well as in the regions which were largely unaffected (e.g., hippocampus, striatum). Our data show that the expression of NRF2 mRNA (*Nfe2l2*) was relatively unaltered between PBS and PFF injected mice (S7A). Remarkably, the expression of NRF2-inhibitor (*Keap1*) was significantly increased in frontal cortex and in the pons region of PFF-injected mice (Fig. 5A), which potentially hints to region specific anti-oxidant response regulation (see Discussion). Among the anti-oxidant genes, the expression levels of three important ROS scavangers, *Hmox1, Gclc and Gsr* (protein: Glutathione S-Reductase- reduces oxidized glutathione disulfide to the anti-oxidant form of glutathione-) were significantly increased in the PFF-injected cohort (Fig. 5B-D). Among other NRF2-related pathways, the changes were either inconsistent (S7B; *Txn*, protein: thioredoxinan anti-oxidant factor in response to intracellular nitric oxide, also inhibits caspase-3 activity), or were not significant (S7C, *Nqo1* and S7D, *PARK7*).

**Figure 5.**
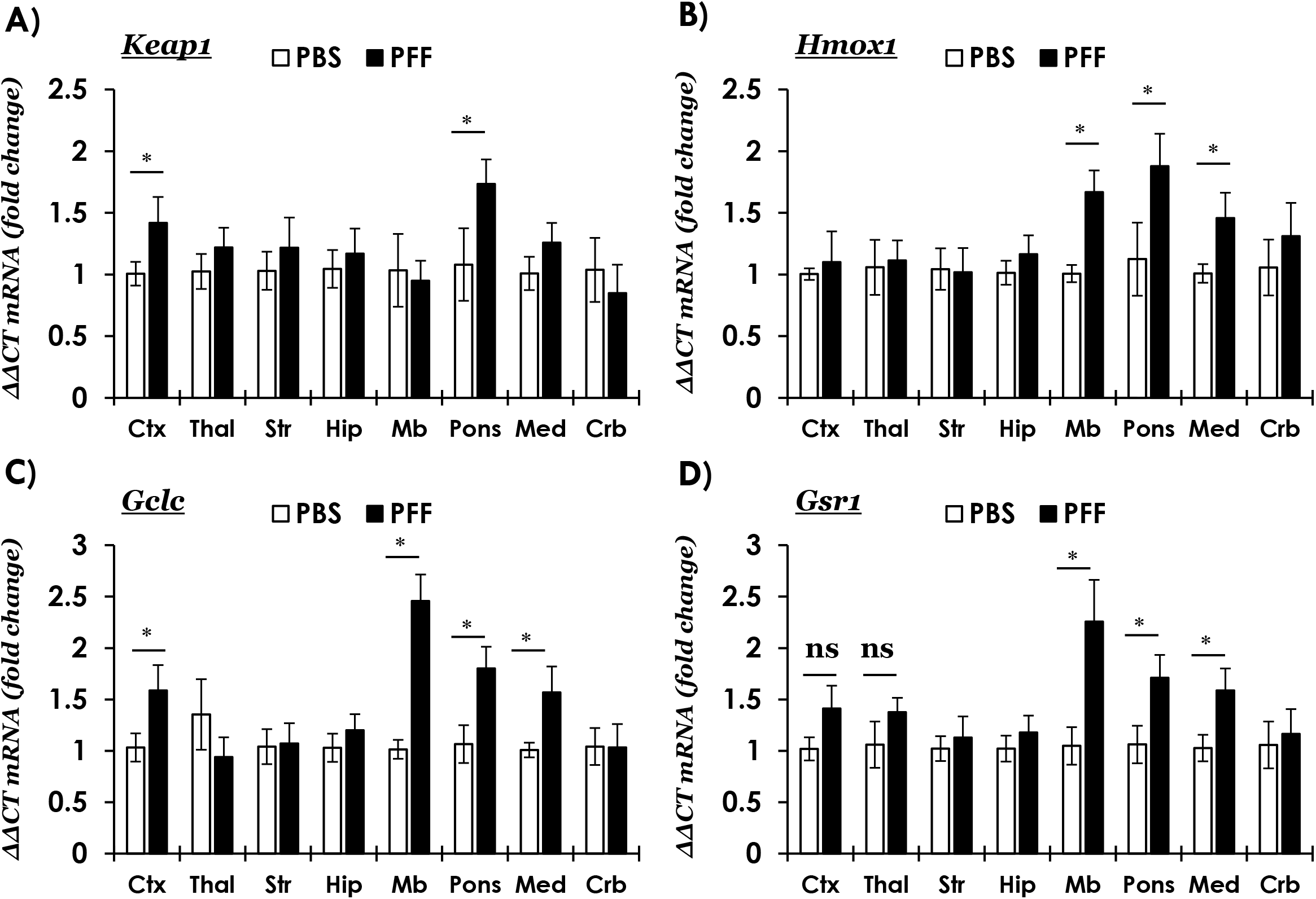
Quantitative RT-PCR analyses of anti-oxidant gene response in the brain regions of PBS or PFF aSyn injected M83^+/+^ mice. **(A)** *Keap1* (Kelch Like ECH Associated Protein 1, NRF2 inhibitor protein), and **(B-D)** NRF2 anti-oxidant response mediators, *Hmox1 (*Heme Oxygenase 1, in B), *Gclc* (Glutamate-Cysteine Ligase Catalytic Subunit, in C) and *Gsr1* (Glutathione-Disulfide Reductase, in D). Relative mRNA quantitation at terminal stage in the following brain regions: frontal cortex (Ctx), thalamus (Thal), corpus striatum (Str), hippocampus (Hip), midbrain (Mb) pons, medulla (Med) and cerebellum (Crb). Mouse *Gapdh* was used as a reference gene (ΔΔCT method: n=6/group, samples were run in duplicates; Error bars represent standard error of the mean, s.e.m.; Pair-wise comparisons were assessed by Mann-Whitney test- only significant differences (*=p≤0.05) are highlighted (ns= not significant); Multiple columns comparisons in One way ANOVA *post-hoc* Bonferroni test were not significant). Also see Figure S7.

In tandem, we also assessed the effects of altered NRF2-anti-oxidant activity on pro-apoptotic and neuroinflammatory pathways, since neuroinflammation has been implicated in PD pathogenesis(18), and is also a characteristic feature in the brains of terminal stage PFF M83 mouse model(36, 47). We found that the expression of mRNAs encoding pro-inflammatory mediators, namely *Tnf-a* (protein, tumor necrosis factor-α) and *Il1-a* (proprotein, interleukin-1α) was significantly elevated in pons and midbrain of PFF-injected M83 mice (S8A-B, respectively). In contrast, the expression of pro-apoptotic effector caspases was relatively unaltered, except localized changes in *Casp3* (S8C, pons) and *Casp6* (S8C, midbrain) in the PFF-injected mice. Hence, the collective tissue response in the brainstem regions- consisting of an increased p NRF2 (S40) nuclear localization (Fig. 4 and S6) and corresponding anti-oxidant gene expression (Fig. 5B-D)- supports the notion that NRF2 homeostatic response is influenced by pathological p-aSyn (p-S129) accumulation.

### Transient aSyn overexpression is not a potent stimulus to activate NRF2 anti-oxidant response in cultured N2A cells

We and others have reported that exogenous application of aggregated aSyn and/or transient aSyn overexpression is associated with altered cellular phenotype including oxidative stress and altered cellular respiration(46, 48, 49). Therefore, we wanted to investigate if acute (short-term) intracellular aSyn accumulation also activates NRF2 anti-oxidant activity and compensatory gene response. Our data show that transient overexpression of WT or mutant human aSyn in cultured neuroblastoma N2A cells did not alter cellular NRF2 expression (Fig. S9A-B), and was associated with slightly increased ROS accumulation (Fig. S9C; compare with hydrogen peroxide-H_2_O_2_ treated groups). In addition, aSyn overexpressing cells did not exhibit a robust anti-oxidant response element (ARE) activity compared with the mock transfected cells (Fig. S9D; also compare with hydrogen peroxide-H_2_O_2_ treated groups). Accordingly, there were non-significant changes in the gene expression, i.e., *Nf2l2, Keap1, Gclc, Gsr1 and Hmox1* (Fig. S9E), which are characteristically activated in this experimental cell model by the application of exogenous ROS inducers(50). Lastly, we detected NRF2 predominantly in diffuse cytoplasmic localization in the cells, with sparse detection of nuclear NRF2 in mock, WT aSyn or A53T aSyn groups (Fig. S9F; compare with S9G- H_2_O_2_ treated groups). These data indicate that the acute (short-term) phase of intracellular aSyn accumulation in this model is not a robust stimulus to activate NRF2, and potentially additional mechanisms may be required for non-physiological aSyn accumulation to mediate these pathways (elaborated in Discussion).

## DISCUSSION

PD represents a multifactorial neurological disorder with underlying role of several genetic risk factors, and is defined by distinct neuropathological subtypes(1, 2, 4). Furthermore, given the complex etiology of PD, it is likely that neuronal dysfunction and demise result from interplay of several factors, including genetic and environmental influences, cellular adaptations to the deleterious effects of mutations increasing disease risk, and neuronal reserve to mitigate metabolic challenges incurred by proteopathic stress(1, 2, 12). Based on the genetics and pathology of PD, significant research efforts have focused on elucidating the consequences of neuronal aSyn accumulation, which could potentially guide the discovery of mechanisms relevant to neurodegeneration(10). In this context, considerable evidence points to the cytotoxic effects of aSyn aggregation, and a prion-like behavior during propagation in the nervous system(4, 8). In particular, several studies point to the deleterious effects of cellular aSyn accumulation on mitochondrial function and impaired energy homeostasis(11, 18). In broader terms, it is plausible that augmenting the activity of cellular mediators promoting mitochondrial function and/or mitigating the deleterious effects of redox imbalance could have a therapeutic potential in PD and related diseases(12, 16).

In this report we show that in PD midbrain showing pathological LB lesions (SN and PAG), there is significant nuclear enrichment of phosphorylated NRF2 (S40), a post-translational modification putatively indicating augmented NRF2 dependent anti-oxidant response (Fig. 1D, 1F and S1B). This is supported by the curated assessment of NRF2-responsive gene expression in independent microarray studies examining disease affected regions in PD (SN, DMX, LC and GPi). Among the examined anti-oxidant genes, analyses of two microarray studies indicate significant changes in the expression of Heme oxygenase, HO-1 (*HMOX1*) in PD SN and GPi (Fig. 2C), which was also reflected in the brainstem of end-stage M83^+/+^ mice (Fig. 5B). These observations are in line with reports showing higher concentrations of HO-1 in the serum of PD patients(51), as well histopathologically found to be enriched in the vicinity of LB inclusions in PD(52). To a considerable extent, data in end-stage M83 mice (Fig. 3-4) implicate pathological aSyn deposition and/or aggregation as a trigger for altered p-NRF2 localization. A tantalizing area for future studies could be to investigate whether misfolded aSyn directly interacts with NRF2, factors controlling NRF2 nuclear accumulation, and/or if such hypothetical interactions affect the NRF2 anti-oxidant response. In this regards, our studies in cultures of dopaminergic N2A cells indicate that ectopic (plasmid) overexpression of aSyn (WT or A53T mutant) was not sufficient to trigger a potent NRF2 nuclear translocation or compensatory gene expression (Fig. S9). These data potentially hint towards the role of aSyn aggregation(48, 49), and the effect of post-translational modifications and/or aSyn truncations characteristically found in disease affected tissues that may alter aSyn protein-protein interactions(10).

Two sets of observations in these data to certain extent are intriguing, and remain largely unexplained: First, the expression profile of the genes examined (e.g., *HMOX1, CASP3*) was not uniformly altered across the PD microarray datasets (Fig. 2, S2-S3). Second, the expression of NRF2 inhibitor was paradoxically increased in the presence of pathological aSyn accumulation, i.e., PD SN (Fig. 2A; GSE7621) and pons of M83 mice (Fig. 5A). With regards to the patient studies, distinct gene expression profiles could reflect heterogeneity in the patient disease stages in the cohorts (e.g., stage of pathology, extent of neurodegeneration, treatment regimen). Importantly, given the putative regulatory role of NRF2 homeostatic response in ROS independent metabolic regulation (e.g., glutamine biogenesis, signal transduction)(23, 31, 53), these data hint towards distinct compensatory mechanism(s) that could be a particular feature of different neuronal populations in a region (e.g., neurotransmitter phenotype, metabolic demands and adaptive response to stress). In addition, cellular differences in inducible NRF2 expression also represents a potential contributor, as reflected by the studies showing that astrocytic NRF2 activation influences the redox homeostasis in neurons, and is a crucial factor in neuroprotection under oxidative stress(38, 54, 55). Lastly, it is also plausible that the increased p-NRF2 (S40) nuclear immunostaining reflects a decreased nuclear export of NRF2 as a result of altered metabolic signaling in additional pathways (e.g., glycogen synthase kinase 3β, Fyn)(31).

From a therapeutic perspective, several small molecules activators of NRF2 pathway restore mitochondrial function under conditions of redox stress, both in cell cultures and also in models of neurodegenerative diseases(26, 28, 56). One such therapeutic molecule, dimethyl fumarate (DMF, Tecfidera), has been successfully used in relapsing-remitting cases of multiple sclerosis, putatively owing to its anti-inflammatory and anti-oxidant effects(57). Furthermore, treatment with DMF, or its active metabolite monomethyl fumarate (MMF), has also been shown to be associated with beneficial outcomes in a number of PD animal models based on (transgenic or viral) aSyn overexpression or chemically induced Parkinsonism(57). These studies show that pharmacological NRF2 activation reduces oxidative stress, improves spinal motor neuron survival, with a concomitant decrease in pathological p-aSyn (S129) accumulation(57). Similarly, increasing NRF2 expression or promoting nuclear localization (by KEAP1 downregulation) rescues the loss of dopaminergic neurons, and improves locomotor performance in PD *Drosophila* models(58, 59). We have previously reported that modulating the activity of a calcium calmodulin eukaryotic elongation factor-2 kinase (eEF2K) exerts a regulatory influence on NRF2 nuclear localization and anti-oxidant response in a ROS-independent manner(50). We showed that eEF2K knockdown improved mitochondrial respiration and ROS scavenging capability of cultured dopaminergic neurons, as well as improved locomotor performance of *C. elegans* expressing human A53T mutant aSyn or human amyloid-β42(46, 50). Notably, we also reported that eEF2K (mRNA) expression and/or activity is pathologically increased in post-mortem PD and Alzheimer disease brains, as well as in relevant transgenic rodent models including the moribund M83^+/+^ mice(46, 50). This highly conserved regulatory pathway plays a crucial role in controlling protein synthesis (an energy demanding process), and also implicated in synaptic function(60, 61). Pharmacological eEF2K inhibition remains an active area of interest for therapeutic discovery in oncology(62); however, the therapeutic potential of this approach in neurodegenerative diseases remains largely untapped.

In conclusion, our data point to a pathological significance of an altered NRF2 response in PD, and warrant further research efforts to assess the potential disease modifying effects of boosting NRF2 in prodromal stages of PD and related diseases.

## MATERIALS AND METHODS

### Human studies

#### Human Tissue processing and Immunohistochemistry (IHC)

Five-micrometer formalin-fixed paraffin embedded post-mortem sections from midbrains of control or PD patients were provided by the laboratory of IM (co-author), as approved by the University of British Columbia Ethics Committee. Anonymized brain sections from 3 control individuals and 5 clinically and pathologically confirmed PD patients were used in these experiments (Table S1).

IHC on brain sections from human tissue was performed after deparaffinization and antigen retrieval. The following antibodies were employed to stain serial tissue sections, as indicated: antibody against phospho-alpha synuclein (p-aSyn, S129; 81A monoclonal; EMD Millipore, #MABN826; dilution 1:1000)(46), and antibody against phospho-NRF2 (p-NRF2, S40; EP1809Y monoclonal; abcam # ab76026); dilution 1:400) using the alkaline phosphatise conjugated streptavidin-biotin ABC kit (Vector Labs, #AK-5000). For destaining/bleaching neuromelanin in substantia nigra in the midbrain sections, the IHC protocol was modified slightly, as described(46). Briefly, sections mounted on slides were incubated in a 60°C degrees oven for 30 minutes and then were transferred into ambient distilled water. Then, the slides were placed in 0.25% potassium permanganate solution for 5 minutes. Subsequently, the slides were rinsed with distilled water. This was followed by incubation in 5% oxalic acid until section became clear. A final rinse in distilled water was performed before proceeding with the normal IHC staining as described above. Sections were counterstained with hematoxylin (Vector Labs, #H-3401). High resolution panoramic images of tissue sections for IHC analyses were acquired using a Leica Aperio digital slide scanner. IHC staining for p-NRF2 (S40) was quantified by manual counting of the DAB (3,3′-diaminobenzidine; Vector Labs, #SK-4100) positive cells.

#### Microarray analyses

Gene expression data from the following trancriptomics datasets was accessed on the Gene Expression Omnibus (GEO)(44) repository of the National Center for Biotechnology Information (NCBI): **1) GSE7621** (*substantia nigra-SN*; Controls, n=9; PD, n=16)(32), **2) GSE43490** (*substantia nigra-SN, dorsal motor nucleus of vagus- DMX and locus coeruleus-LC*; Controls, n=5-7; PD, n=8)(33), **3) GSE20146** (*globus pallidus, interna*-*GPi*; Controls, n=10; PD, n=10)(35) and **4) GSE26927** (*substantia nigra-SN*; Controls, n=7; PD, n=12)(34). Unique probe identities for the transcripts, and additional microarray platform information is provided in Table S2.

### Animal studies

#### Animal husbandry

Transgenic M83 mice [B6;C3-Tg(Prnp-SNCA*A53T)83Vle/J](37) were housed at the Bartholin animal facility at Aarhus University in accordance with Danish regulations and the European Communities Council Directive for laboratory animals, under the authorization #2017-15-0201-01203 issued to PHJ (co-author). The mice we housed under 12 hours light/dark cycle and fed with regular chow diet *ad libitum*. The experiments were performed using both male and female mice.

#### Intramuscular injection of preformed aSyn fibrils

Mouse aSyn fibrils were prepared and characterized *in vitro* (for purity, biophysical properties and biological activity) essentially as described(45). Homozygous M83^+/+^ mice (2-3 months old, n=8/group) were bilaterally inoculated with a single injection (5 μl) of recombinant mouse aSyn preformed fibrils (PFF, 2 mg/mL in phosphate buffered saline- PBS)- or PBS vehicle- into the hindlimb biceps femoris (using a 10-μL Hamilton syringe with a 25-gauge needle) under isoflurane (1-5%) anesthesia(45, 46). Separate syringes were used for each type of inoculums (PBS or PFF) to avoid cross-contamination. After the injection, the mice were allowed to recover and returned to their original housing cages.

#### Tissue collection

Mice were euthanized with an overdose of sodium pentobarbital (150 mg/kg, intraperitoneal) and perfused with ice-cold PBS pH 7.4 containing phosphatase inhibitors (25 mM β-glycerolphosphate, 5 mM NaF, 1 mM Na3VO4, 10 mM Na-pyrophosphate). Brains were collected and one hemisphere was processed for IHC (see below). The contralateral hemisphere was further microdissected under the microscope to isolate the brain region of interest on ice-cold sterile filtered PBS supplemented with 10 mM D- glucose (ThermoFisher # A2494001), snap frozen in liquid nitrogen, and stored at −80°C for total RNA extraction (see below).

#### IHC

IHC on 10μm thick sections from formalin fixed paraffin embedded tissue was performed after deparaffinization and antigen retrieval in citrate buffer pH-6.0. Following primary antibodies were employed: phospho-αSyn-S129 (11A5 monoclonal, kind gift to PHJ by Imago Pharmaceuticals- 1:1000)(45) and phospho-NRF2-S40 (rabbit polyclonal, ThermoFisher #PA5-67520-; 1:500). For IHC, DAB (Electron Microscopy Sciences #13082) chromogen detection was performed following incubation with biotin conjugated secondary antibodies and Extra-Avidin peroxidise (Sigma #E2886- 1:200). Sections were counterstained with hematoxylin (Vector Labs, #H-3401). High resolution views were obtained using Olympus VS120 digital slide scanner and 5-40X views were extracted using OlyVia software (Olympus). Panoramic digital slide scans were mapped onto Mouse Brain Atlas to neuroanatomically define the regions/nuclei (*Paxinos and Franklin’s The Mouse Brain in Stereotaxic Coordinates*, *Elsevier Publishing, 4^th^ Ed*.)(63).

#### Quantitative RT-PCR

Total RNA was extracted from the fresh frozen tissue using QIAzol lysis reagent (Qiagen, #74134) and purified using a commercial kit (Qiagen, #74134). cDNA was synthesized from 500ng of total RNA using high capacity reverse transcriptase kit (Applied Biosystems, #4368814). RT-qPCR was performed using SYBR green (Thermo Fisher, #4385616) under standard conditions with unique primer pairs (Table S3) in duplicate samples. The data were analyzed by relative ΔΔCT quantification method using the murine glyceraldehyde 3-phosphate dehydrogenase (*Gapdh)* cycle (CT) values as the internal reference in each sample(64).

## Cell culture

### aSyn plasmid transient overexpression and Nrf2 anti-oxidant response in cultured N2A cultures

Mouse neuroblastoma (N2A) cells were obtained from ATCC (#CCL-131), and maintained in DMEM (4.5 g/L glucose; Gibco, #11965-084) supplemented with 1% antibiotic-antimycotic solution (Gibco, #15240062) and 10% Fetal Bovine Serum (FBS). The cells were cultured in 6-well (500, 000 cells/well) 12-well (250,000 cells/well) or 96-well (50,000 cells/well) plates. DNA plasmid transfections were performed using Lipofectamine 2000 (Invitrogen, #11668019) according to the recommended procedures. After 24 hours, cells were briefly washed with phosphate-buffer saline (PBS) and allowed to differentiate into neurons in a modified culture medium containing DMEM (Gibco, #21969035) supplemented with 500 μM Lglutamine, 1% antibiotic-antimycotic, 2% FBS and 500 μM Dibutyryladenosine 3′,5′-cyclic monophosphate (db cAMP; Sigma, #D0627)(46). Unless indicated otherwise, differentiated N2A cells which were mock transfected, or transfected with AS plasmids (ASyn-WT or ASyn-A53T; Addgene plasmid #40824 and #40825 respectively) were used in the assays after 72-76 hours post-transfection. Cellular ROS measurements were performed after incubation with 5 μg/mL 2,7-dichlorofluorescein diacetate- H2DCFDA (ThermoFisher #D399) for 30 minutes in fresh medium (± pretreatment with hydrogen peroxide, H_2_O_2_; 250 μM for 2 hours) according to the manufacturer’s protocol, and essentially as described(46, 50). Anti-oxidant response element (ARE, Nrf2) promoter activity was assessed by ARE Reporter kit (BPS Biosciences #60514) combined with dual luciferase assay (Promega #E1910), (± pretreatment with hydrogen peroxide, H_2_O_2_; 250 μM for 2 hours)(50). Gene expression analyses were performed using identical primer pairs (Table S3) and experimental parameters as described under the mouse tissues above. Immunoflourescence microscopy (± pretreatment with hydrogen peroxide, H_2_O_2_; 250 μM for 2 hours) was performed after gentle fixation (4% PFA, 10 min, 4°C) followed by incubation with rabbit polyclonal NRF2 antibody (Novus Bioogicals #NBP1-32822, 1:100) and Alexa488 fluorophore conjugated goat secondary antibody (ThermoFisher #A-11008). Cell nuclei were labelled with fluorescent DNA marker DRAQ5 (Biostatus # DR50050) and images were acquired acquired using a Zeiss observer inverted microscope equipped with colibri 7 LED illumination, and operated using Zen (Zeiss) software.

## Statistics

The data were statistically analyzed in Graphpad Prism software (version 9) and final graphs were prepared in Microsoft Excel. Data were analyzed by One-Way ANOVA, and pair-wise comparisons were statistically assessed by the Mann-Whitney nonparametric test or student’s ttest as indicated in the respective figure legends.

## Supporting information

Delaidelli A et al SI

## DECLARATIONS, AS APPLICABLE

### Availability of data and material

The transcriptomic datasets analyzed during this study can be accessed on the NCBI GEO webpage(44), with the accession and probe IDs provided in Table S2. Otherwise, all the data generated and analyzed during this study are included in the main manuscript or the associated supplementary files.

## Acknowledgements

The authors would like to thank Helene Andersen (JRN lab) and Sandra Bonnesen (CV) for technical help during the study.

