## Supplementary material for "α-Synuclein pathology in Parkinson disease activates homeostatic NRF2 anti-oxidant response": Delaidelli A et al SI

**Author Affiliations:** [¶]Danish Research Institute of Translational Neuroscience (DANDRITE)-Nordic-EMBL Partnership for Molecular Medicine, Department of Biomedicine, Aarhus University, DK-8000, Aarhus C, Denmark; [†]Core Center for Molecular Morphology, Section for Stereology and Microscopy Department of Clinical Medicine, Department of Pathology, Aarhus University Hospital, DK-8200 Aarhus N, Denmark; [ǂ]Department of Pathology and Laboratory Medicine, University of British Columbia, Vancouver, Canada; [‡]British Columbia Cancer Research Centre, Vancouver, BC V5Z 1L3 Canada.

[§]**Address correspondence to**:

**Contents:**

- **Supplementary Figures (S1-S9) and figure legends**
- **Tables S1-S3**

**Supplementary Figures**

**S1. Immunostaining of phospho-aSyn (S129) and phospho-NRF2 (S40) in postrmoretm midbrain sections of a control and a PD case**. **(A-B)** Representative IHC images showing panoramic and 20X magnified views of phospho-aSyn (S129) and phospho-NRF2 (S40) immunostaining in midbrain (PAG, periaqueductal grey and SN, substantia nigra) sections of a control (in A) and a PD case (in B). Red arrows in the insets point to cells with predominant nuclear localization of phospho-NRF2 (S40) (scale bar in 20X views=100 µm). Also see Figure 1 (data from additional controls and PD cases- Table S1). Primary antibodies: p-aSyn (S129)- 81A and p-NRF2 (S40)- EP1809Y.

**S2. Curated gene expression analyses of publicly available microarray datasets in PD using Gene Expression Omnibus (GEO).** **(A-D)** Anti-oxidants *GSR1* (Glutathione-Disulfide Reductase, in A), *PARK7* (Parkinsonism Associated Deglycase- DJ1, in B), *NQO1* (NAD(P)H Quinone Dehydrogenase 1, in C) and *TXN* (Thioredoxin, in D). **(E-F)** Inflammatory mediators, *TNFA* (Tumor Necrosis Factor alpha, in C) and *IL1A* (Interleukin 1, in D). The values across the datasets are expressed relative to the controls in each microarray dataset, i.e., mean value of control samples=1 (a.u., arbitrary units). Error bars represent standard deviation of the mean, s.d. In GSE7621- SN (*substantia nigra*); controls (Ctrl, n=9) and PD (Parkinson Disease cases, n=16); in GSE43490- SN (*substantia nigra*; controls, n=6 and PD, n=8), DMX (*dorsal motor nucleus of vagus*; controls, n=5 and PD, n=8), LC (*locus coeruleus*; controls, n=7 and PD, n=8); in GSE20146- GPi (*globus pallidus interna*; controls, n=10 and PD, n=10). Pair-wise comparisons were assessed by Mann-Whitney test- only significant differences (*=p≤0.05, **=p≤0.01) are highlighted. Probe IDs, microarray platforms and relevant studies are listed in Table S2. Also see Figure 2.

**S3. Curated gene expression analyses of a publicly available microarray dataset (GSE26927) in selected neurodegenerative diseases using Gene Expression Omnibus (GEO). (A)** *NFE2L2* (Nuclear Factor, Erythroid 2 Like 2, NRF2)**, (B)** *KEAP1* (Kelch Like ECH Associated Protein 1, NRF2 inhibitor protein), **(C-F)** Anti-oxidants *GSR1* (Glutathione-Disulfide Reductase, in C), *GCLC* (Glutamate-Cysteine Ligase Catalytic Subunit, in D), *TXN* (Thioredoxin, in E) and *PARK7* (Parkinsonism Associated Deglycase- DJ1, in F). Disease conditions include: PD (Parkinson Disease), AD (Alzheimer Disease), HD (Huntington Disease), ALS (Amyotrophic Lateral Sclerosis), MS (Multiple Sclerosis). Samples include: *substantia nigra* (controls n=7, PD n=12), *entorhinal cortex* (controls n=6, AD n=12), *cervical spinal cord* (controls n=7, ALS n=9), *nucleus caudatus* (controls n=10, HD n=10), and *grey matter lesions in frontal gyri* (controls n=10, MS n=10). Error bars represent standard deviation of the mean, s.d. Pair-wise comparisons were assessed by Mann-Whitney test- only significant differences (*=p≤0.05, **=p≤0.01, ***=p≤0.001) are highlighted. Probe IDs, microarray platform and relevant study are listed in Table S2.

**S4. Immunostaining of phospho-aSyn (S129) in the brain regions of PBS injected M83$^{+/+}$ mice. (A)** Representative IHC images showing phospho-aSyn (S129) immunostaining (10X low magnification views and 40X magnified views in the insets; scale bar=200 µm; Aq. in the image showing PAG, cerebral aqueduct; 4v. in the image showing vestibular nuclei image, 4$^{th}$ ventricle). Bregma co-ordinates for the brain regions were determined according to Paxinos and Franklin: (Bregma, -4.03 mm) midbrain at the level of superior colliculi showing PAG, red nucleus and tegmentum; (Bregma, -5.91) pontocerebellar junction showing cerebellar nuclei, cerebellar lobules (*cb1-5*), vestibular nuclei and pontine gigantocellualr nuclei (Gi); and (Bregma, 0.49 mm) forebrain showing motor cortex (M1 and M2) and corpus striatum). Also see Figure 3 (p-aSyn, S129 in brain regions of PFF aSyn injected M83$^{+/+}$ mice). Primary antibody in A: p-aSyn (S129)- 11A5.

**S5. Immunostaining of phospho-NRF2 (S40) in the brain regions of PBS injected M83$^{+/+}$ mice. (A)** Representative IHC images showing phospho-NRF2 (S40) immunostaining. Notice variable degrees of staining, prominently cytoplasmic, detected in red nucleus, midbrain tegmentum, motor cortex and corpus striatum (10X low magnification views and 40X magnified views in the insets; scale bar=200 µm; Aq. in the image showing PAG, cerebral aqueduct; 4v. in the image showing vestibular nuclei image, 4$^{th}$ ventricle). Bregma co-ordinates for the brain regions were determined, according to Paxinos and Franklin: (Bregma, -3.79 mm) midbrain at the level of superior colliculi showing PAG, red nucleus and tegmentum; (Bregma, -5.79) pontocerebellar junction showing cerebellar nuclei, cerebellar lobules (*cb1-5*), vestibular nuclei and pontine gigantocellualr nuclei (Gi); and (Bregma, 0.73 mm) forebrain showing motor cortex (M1 and M2) and corpus striatum. Also see Figure 4 (p-NRF2, S40 in brain regions of PFF aSyn injected M83$^{+/+}$ mice). Primary antibody in A: p-NRF2 (S40)- PA5-67520.

**S6. Immunostaining of phospho-aSyn (S129) and phospho-NRF2 (S40) in select brain regions of M83$^{+/+}$ mice. (A-B)** Representative IHC 40X magnified views showing phospho-aSyn (S129) in (A) and phospho-NRF2 (S40) immunostaining in (B), in the select brain regions of three additional animals from PFF injected cohort. Primary antibodies: p-aSyn (S129)- 11A5 and p-NRF2 (S40)- PA5-67520. **(C)** Phospho-NRF2 (S40) immunostaining in select brain regions of three additional animals from the PBS injected cohort. The animals in each cohort are designated #1, #2 and #3. Regions examined include: periaqueductal grey (PAG), red nucleus (RN), mesencephalic tegmentum (Teg.), pontine gigantocellular nuclei (Gi), pontine vestibular nuclei (VN) and deep cerebellar nuclei (DCN). Primary antibody in C: p-NRF2 (S40)- PA5-67520. Scale bar in A-C=25 µm. Also see Fig. 3-4 and S4-5.

**S7. Quantitative RT-PCR analyses of anti-oxidant gene response in the brain regions of PBS or PFF aSyn injected M83$^{+/+}$ mice. (A)** *Nfe2l2* (Nuclear Factor, Erythroid 2 Like 2, NRF2), and **(B-D)** NRF2 anti-oxidant response mediators, *Txn* (Thioredoxin, in B), *Nqo1* (NAD(P)H Quinone

Dehydrogenase 1, in C) and *Park7* (Parkinsonism Associated Deglycase- DJ1, in D). Relative mRNA quantitation at terminal stage in the following brain regions: frontal cortex (Ctx), thalamus (Thal), corpus striatum (Str), hippocampus (Hip), midbrain (Mb) pons, medulla (Med) and cerebellum (Crb). Mouse *Gapdh* was used as a reference gene (ΔΔCT method: n=6/group, samples were run in duplicates; Error bars represent standard error of the mean, s.e.m.; Pair-wise comparisons were assessed by Mann-Whitney test- only significant differences (*=p≤0.05) are highlighted (ns= not significant); Multiple columns comparisons in One way ANOVA *post-hoc* Bonferroni test, were not significant). Also see Figure 5.

**S8. Quantitative RT-PCR analyses of inflammatory gene response in the brain regions of PBS or PFF aSyn injected M83$^{+/+}$ mice. (A-B)** Inflammatory mediators, *Tnfa* (Tumor Necrosis Factor alpha, in A) and *Il1a* (Interleukin 1, in B). **(C-D)** Pro-apoptotic caspases**,** *Casp3* (Caspase 3, in C) and *Casp6* (Caspase 6, in D). Relative mRNA quantitation at terminal stage in the following brain regions: frontal cortex (Ctx), thalamus (Thal), corpus striatum (Str), hippocampus (Hip), midbrain (Mb) pons, medulla (Med) and cerebellum (Crb). Mouse *Gapdh* was used as a reference gene (ΔΔCT method: n=6/group, samples were run in duplicates; Error bars represent standard error of the mean, s.e.m.; Pair-wise comparisons were assessed by Mann-Whitney test- only significant differences (*=p≤0.05) are highlighted (ns= not significant); Multiple columns comparisons in One way ANOVA *post-hoc* Bonferroni test, were not significant).

**S9. Effects of transient aSyn overexpression on NRF2 anti-oxidant response in cultured N2A cells. (A-B)** Western immunoblotting showing the expression of aSyn (WT-wild type and mutant A53T aSyn and Nrf2 (in A) and corresponding densitometry analyses (in B). Antibodies used in A: aSyn (BD Biosciences, clone 42 #610787) and Nrf2 (Novus Biological #NBP1-32822). Expression values in S9B were normalized using β- actin as the loading control. The error bars indicate Mean ± S.D. (n=6 biological replicates/group from 3 independent experiments). **(C)** ROS measurements by $H_2$-DCFDA assay in mock transfected cells or cells expressing aSyn (WT and A53T), following treatment with PBS or exposure to exogenous $H_2O_2$ (250 μM; 2 hours). Fluorescence values were normalized to the control cells (mock transfected, PBS treated; Mean expressed as 100%). The error bars indicate Mean ± s.e.m. (n=9-12 biological replicates/group; Pair-wise comparisons, t-test, **=0.01, ***=0.005) from 3-4 independent experiments). **(D)** Nrf2 ARE promoter activity in mock transfected cells or cells expressing aSyn (WT and A53T), following treatment with PBS or exposure to exogenous $H_2O_2$ (250 μM; 2 hours). Luminescence values were normalized to the control cells (mock transfected, PBS treated; Mean expressed as 1.0; a.u.=arbitrary units). The error bars indicate Mean ± s.e.m. (n=8 biological replicates/group; Pair-wise comparisons, t-test, ***=0.005 from 3 independent experiments). **(E)** Relative (ΔΔCT) expression of cytoprotective anti-oxidant genes in mock transfected cells or cells expressing aSyn (WT and A53T): *Nfe2l2, Keap1, Gclc, Gsr1 and Hmox1*. The data are representative of 3 independent experiments. The error bars indicate Mean ± S.D. (n=2 biological replicates/group, run in duplicates in each experiment; Pair-wise comparisons in t-test were not significant). **(F-G)** Immunofluorescence (IF) microscopy analyses of Nrf2 (antibody, NBP1-32822) in mock transfected cells or cells expressing aSyn (WT and A53T), following treatment with PBS (in F) or exposure to exogenous $H_2O_2$ (in G; $H_2O_2$ treatment: 250 μM for 2 hours). DRAQ5 was used as a nuclear marker. Yellow arrows indicate select instances of Nrf2 IF detection in the cell nucleus (scale bar= 50 μm).

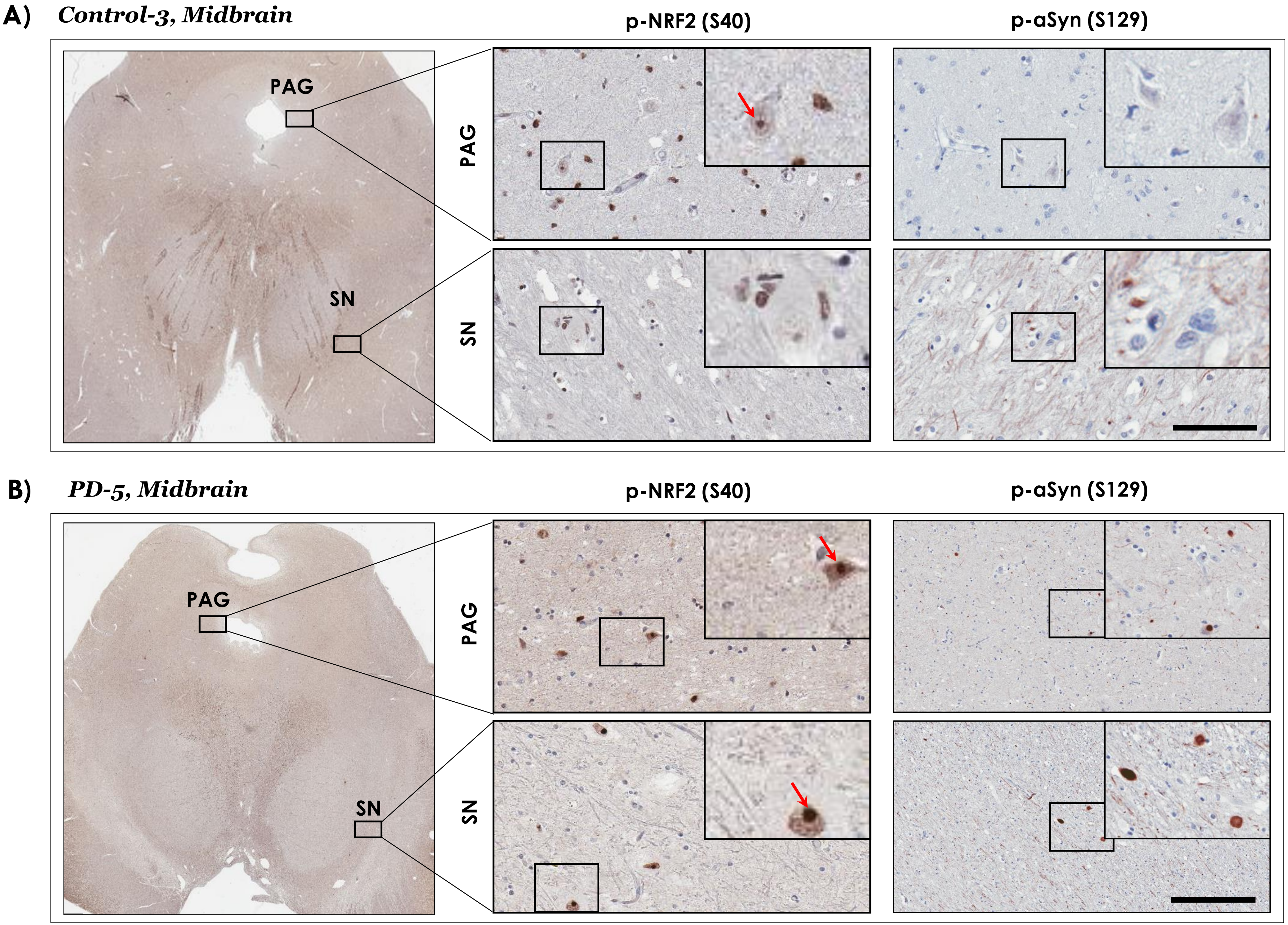
A)
Control-3, Midbrain
p-NRF2 (S40)
p-aSyn (S129)
PAG
SN
PAG
SN
B)
PD-5, Midbrain
p-NRF2 (S40)
p-aSyn (S129)
PAG
SN
PAG
SN

Fig. S2

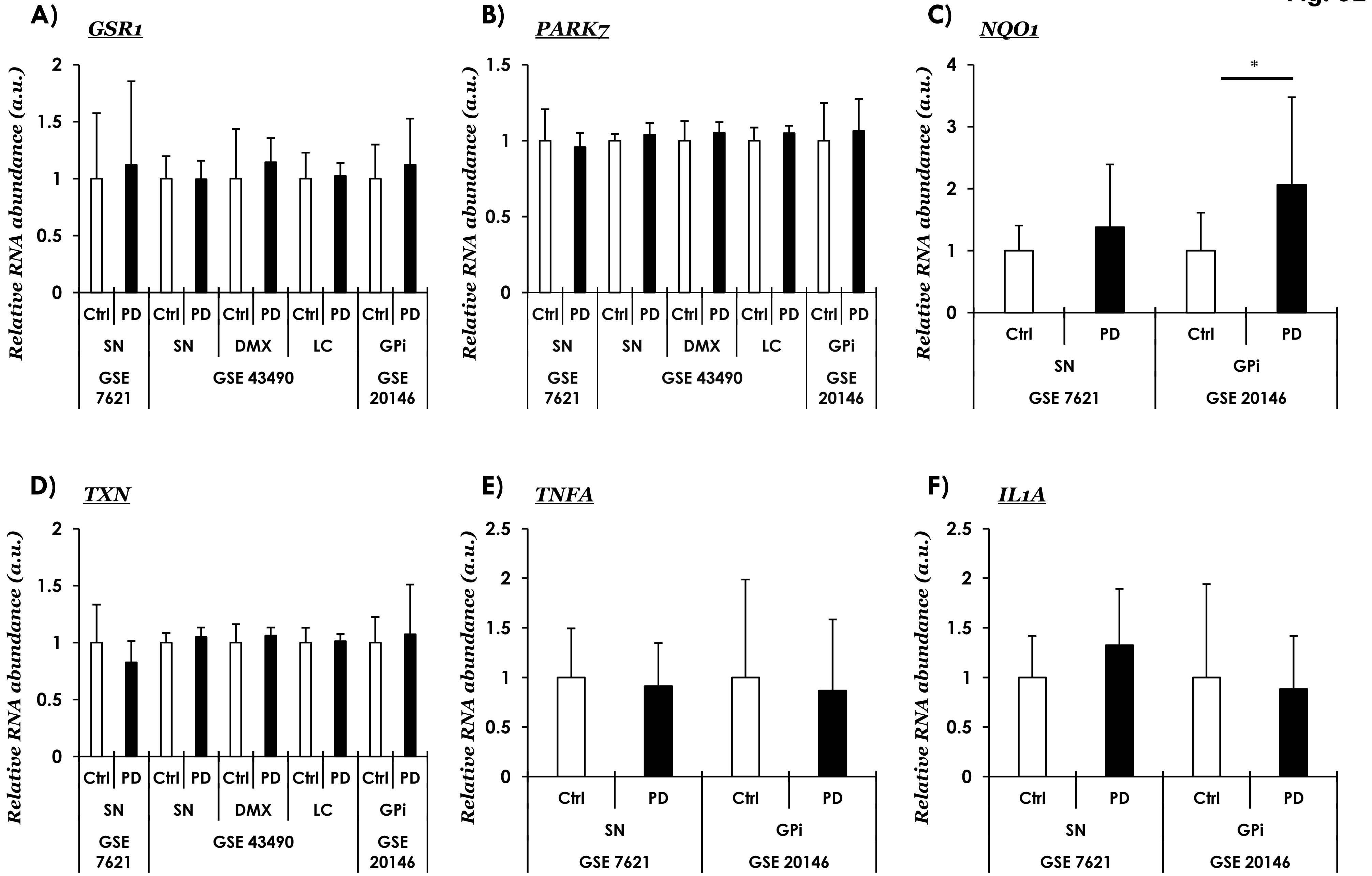

A)
GSR1
Relative RNA abundance (a.u.)
2
1.5
1
0.5
0
Ctrl
PD
SN
DMX
LC
GPi
GSE 7621
GSE 43490
GSE 20146
B)
PARK7
1.5
1
0.5
0
C)
NQO1
4
3
2
1
0
*
D)
TXN
E)
TNFA
2.5
F)
IL1A

### GSE26927

Fig. S3

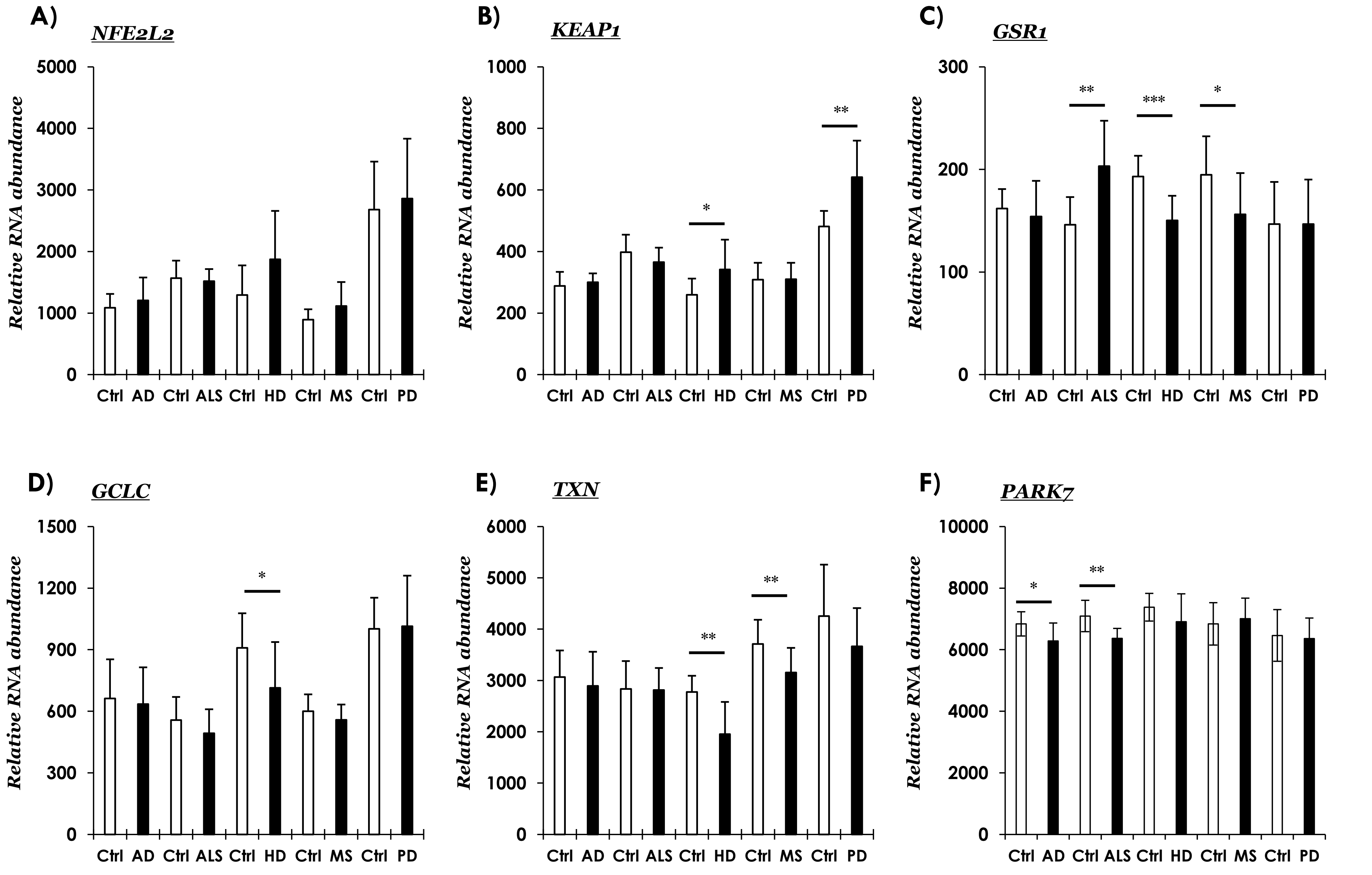

A) *PBS injected* p-aSyn (S129) Fig. S4

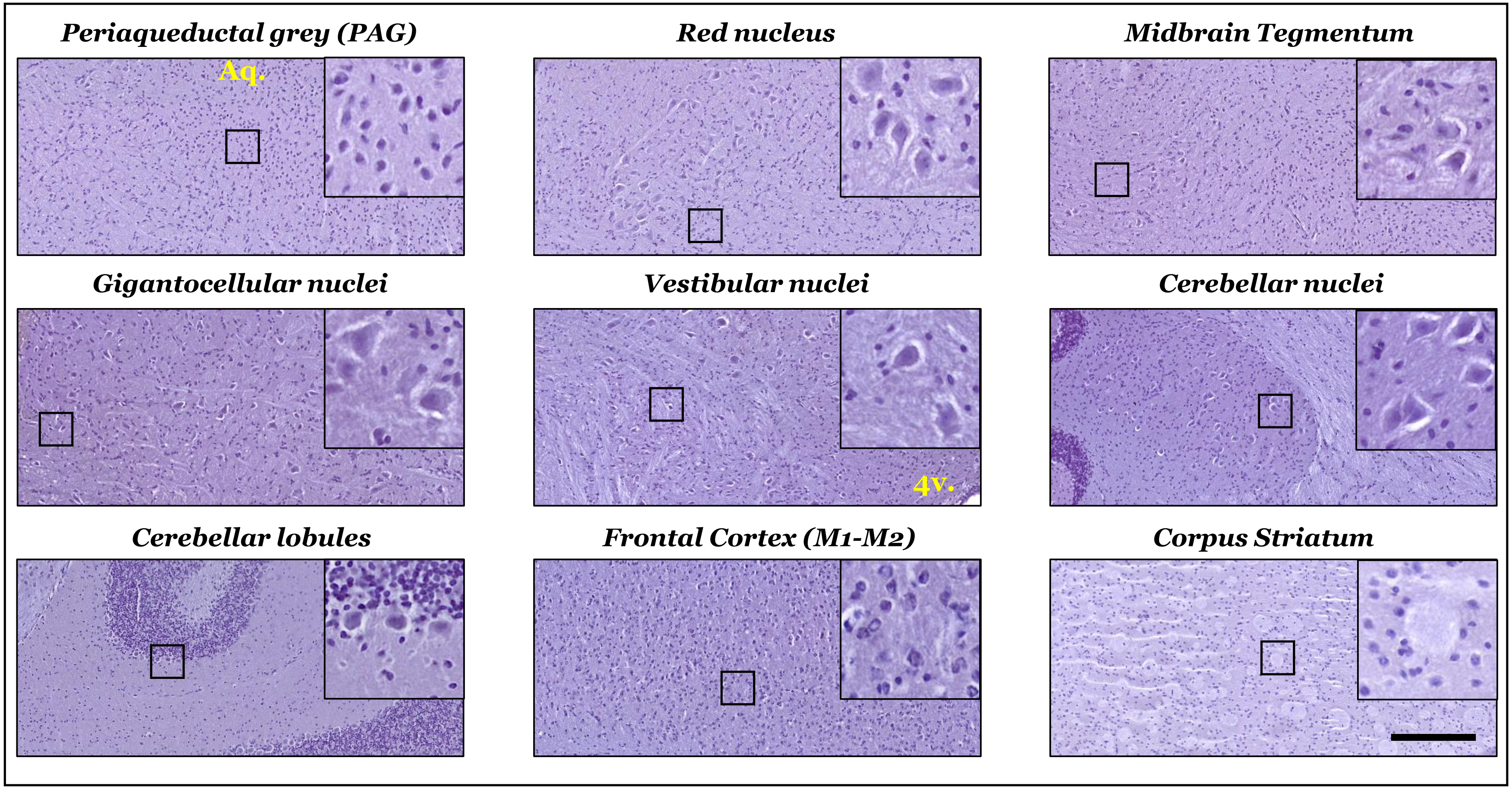

Fig. S5

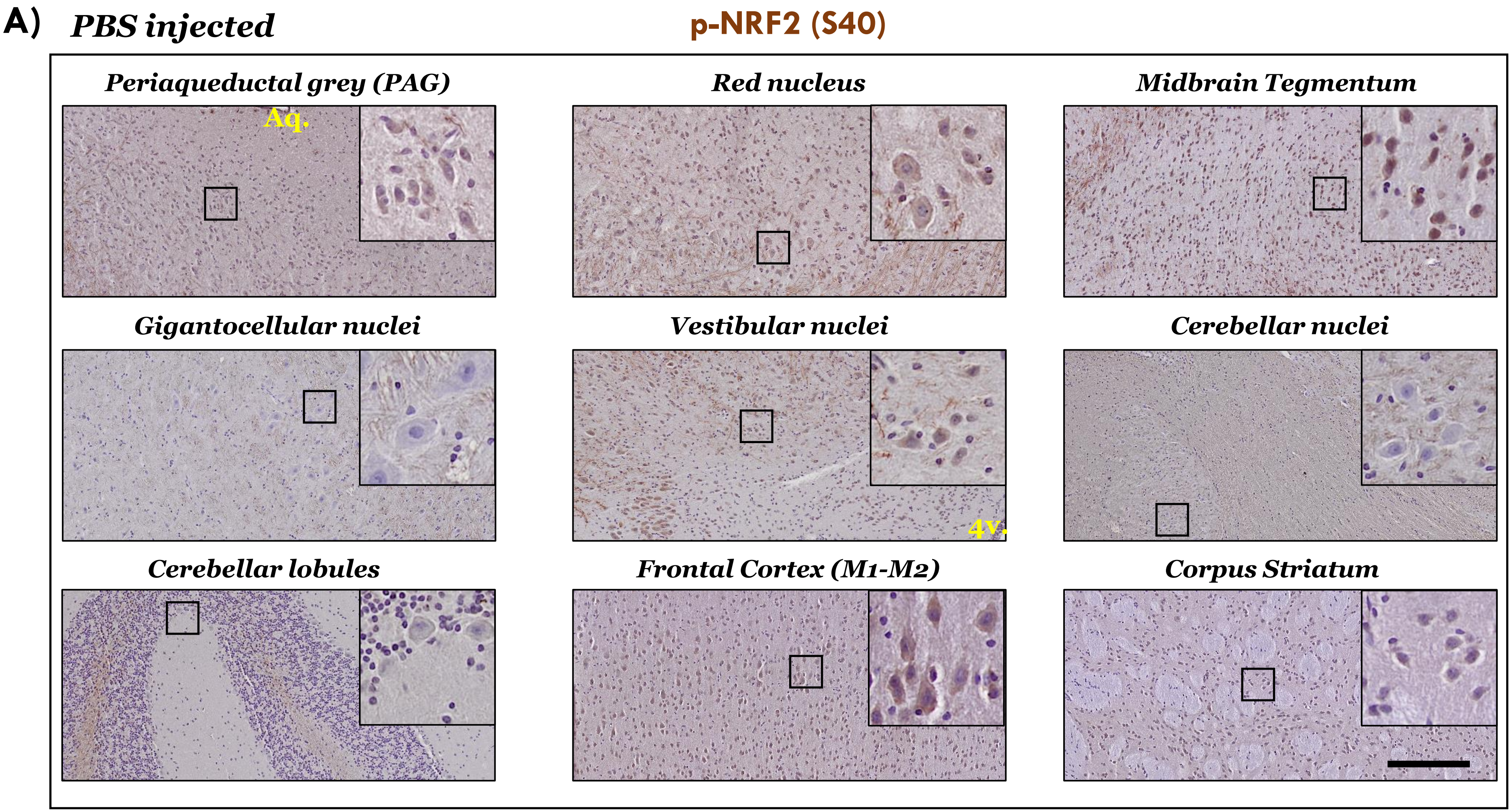
A)
PBS injected
p-NRF2 (S40)
Periaqueductal grey (PAG)
Aq.
Red nucleus
Midbrain Tegmentum
Gigantocellular nuclei
Vestibular nuclei
4v.
Cerebellar nuclei
Cerebellar lobules
Frontal Cortex (M1-M2)
Corpus Striatum

Fig. S6

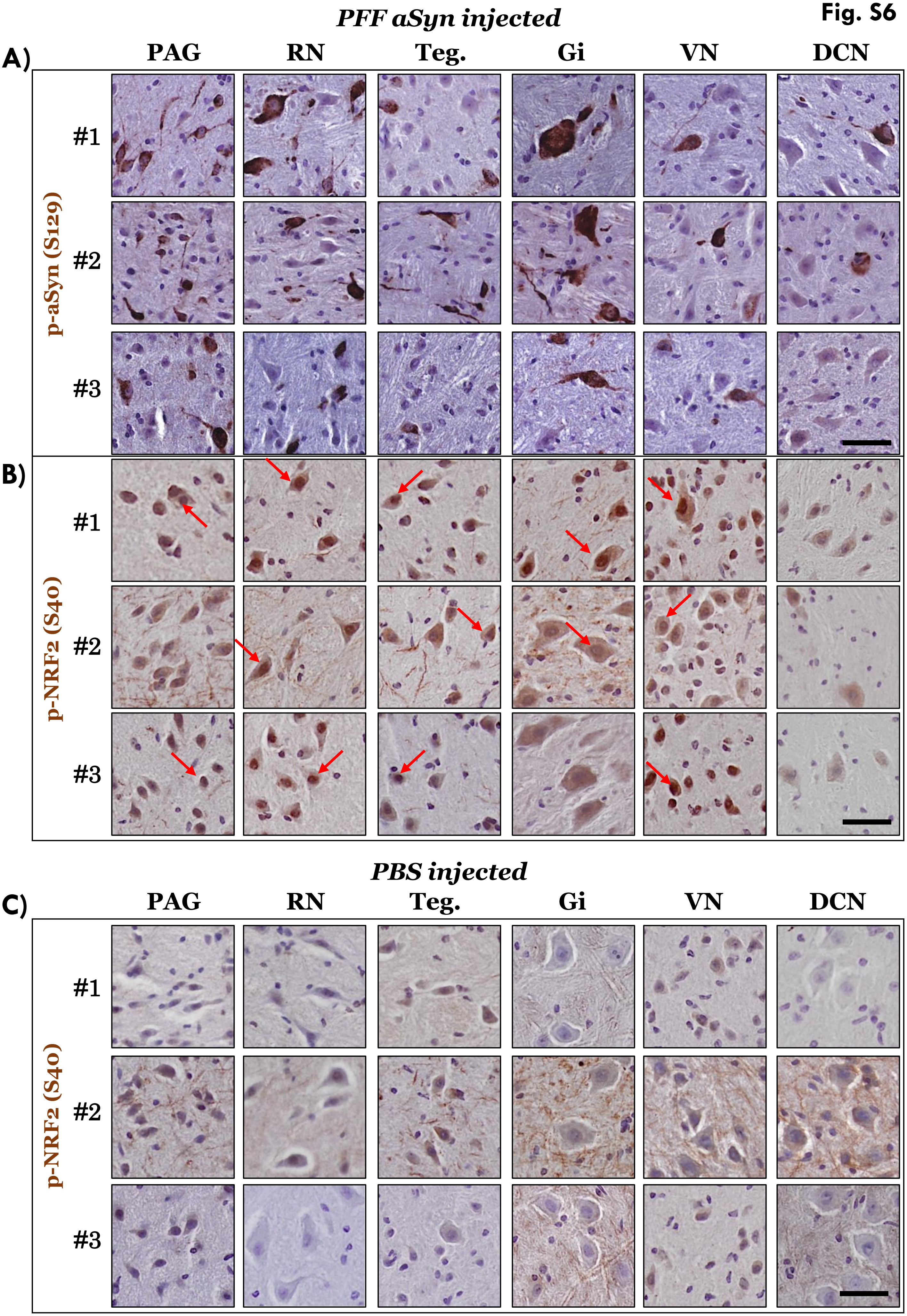

PFF aSyn injected
A)
PAG
RN
Teg.
Gi
VN
DCN
p-aSyn (S129)
#1
#2
#3
B)
p-NRF2 (S40)
#1
#2
#3
PBS injected
C)
PAG
RN
Teg.
Gi
VN
DCN
p-NRF2 (S40)
#1
#2
#3

Fig. S7

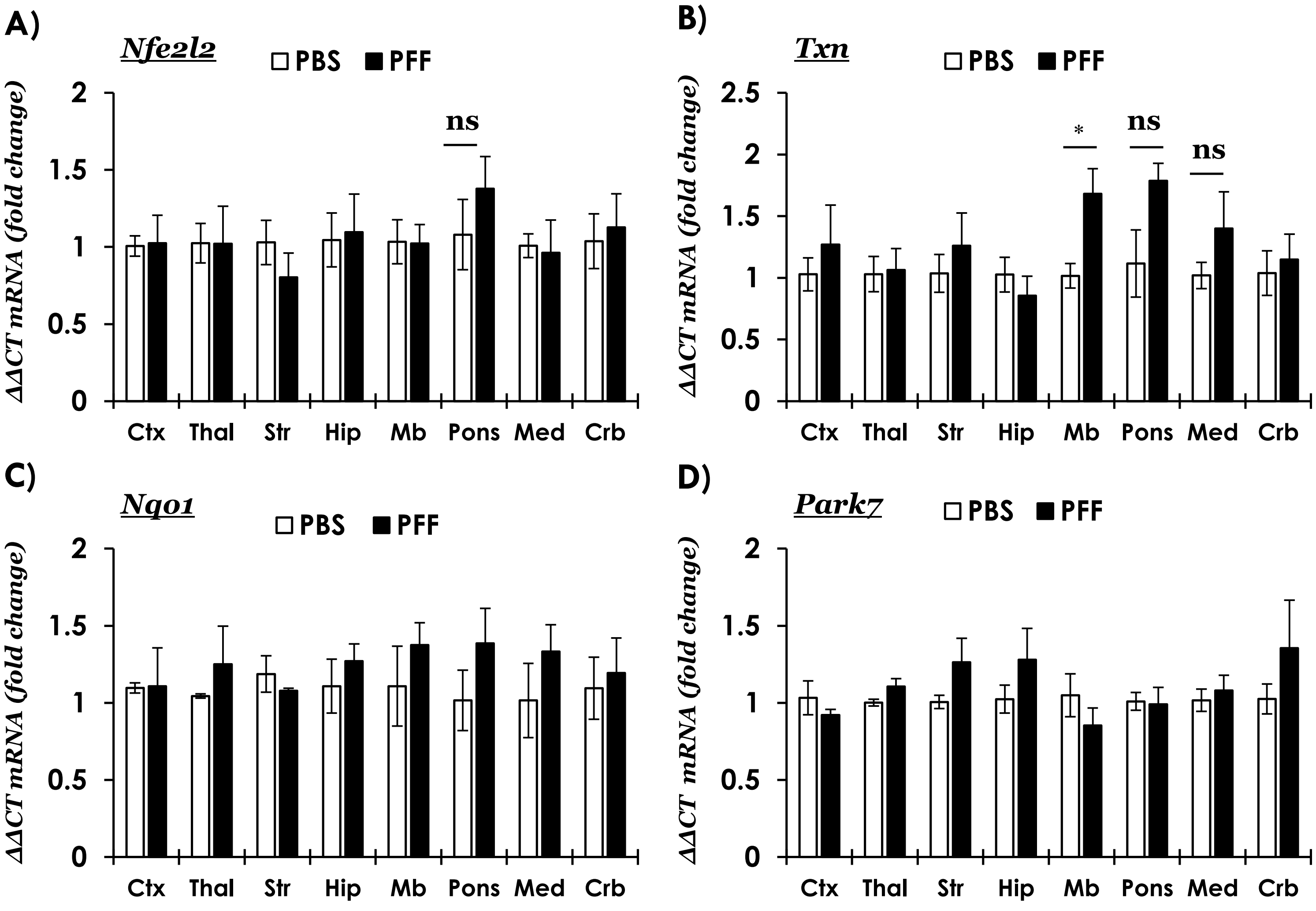
A)
Nfe2l2
PBS
PFF
ΔΔCT mRNA (fold change)
2
1.5
1
0.5
0
ns
Ctx
Thal
Str
Hip
Mb
Pons
Med
Crb
B)
Txn
PBS
PFF
ΔΔCT mRNA (fold change)
2.5
2
1.5
1
0.5
0
*
ns
ns
Ctx
Thal
Str
Hip
Mb
Pons
Med
Crb
C)
Nqo1
PBS
PFF
ΔΔCT mRNA (fold change)
2
1.5
1
0.5
0
Ctx
Thal
Str
Hip
Mb
Pons
Med
Crb
D)
Park7
PBS
PFF
ΔΔCT mRNA (fold change)
2
1.5
1
0.5
0
Ctx
Thal
Str
Hip
Mb
Pons
Med
Crb

Fig. S8

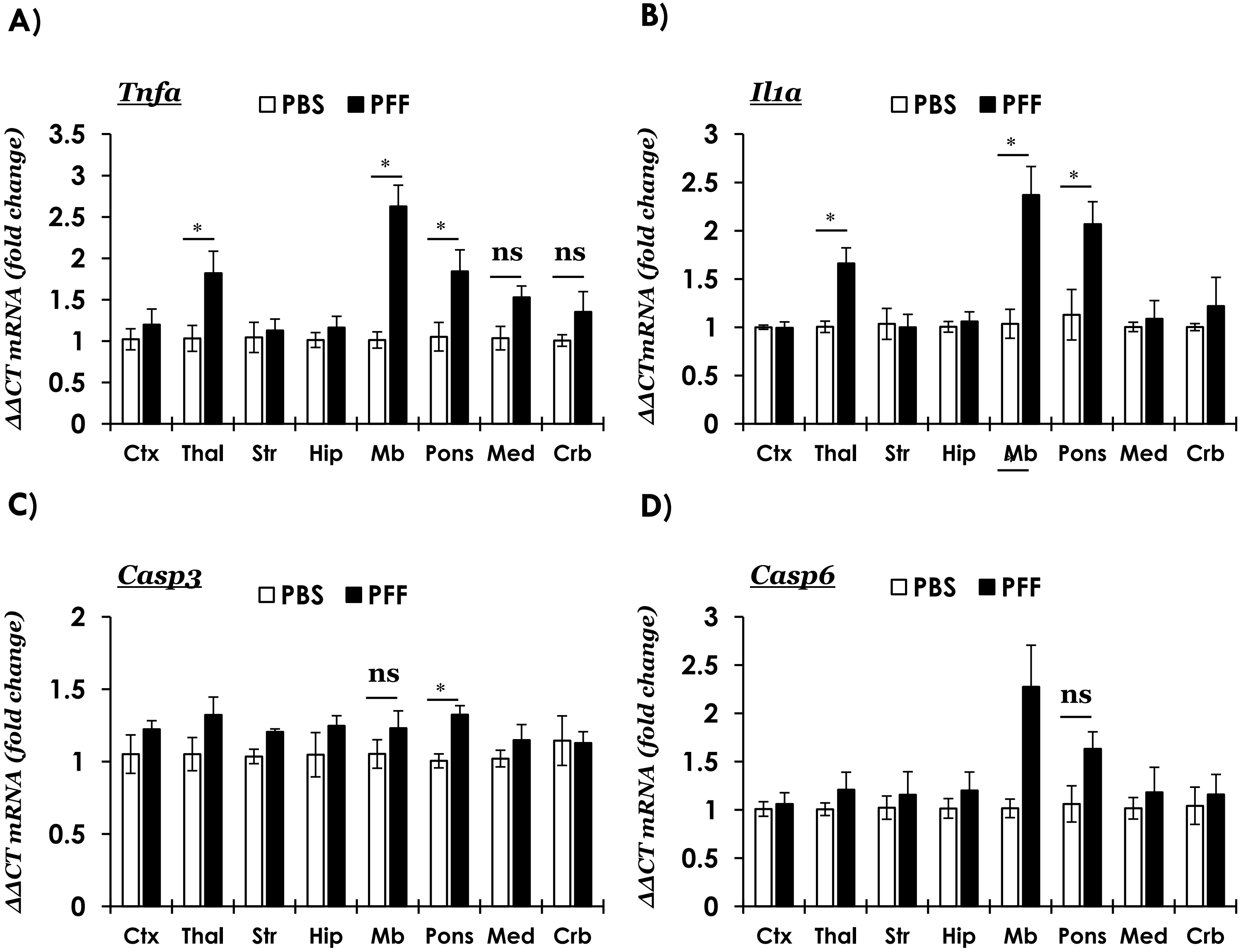
A)
Tnfa
PBS
PFF
ΔΔCT mRNA (fold change)
3.5
3
2.5
2
1.5
1
0.5
0
*
*
*
ns
ns
Ctx
Thal
Str
Hip
Mb
Pons
Med
Crb
B)
Il1a
PBS
PFF
ΔΔCT mRNA (fold change)
3
2.5
2
1.5
1
0.5
0
*
*
*
Ctx
Thal
Str
Hip
Mb
Pons
Med
Crb
C)
Casp3
PBS
PFF
ΔΔCT mRNA (fold change)
2
1.5
1
0.5
0
ns
*
Ctx
Thal
Str
Hip
Mb
Pons
Med
Crb
D)
Casp6
PBS
PFF
ΔΔCT mRNA (fold change)
3
2.5
2
1.5
1
0.5
0
ns
Ctx
Thal
Str
Hip
Mb
Pons
Med
Crb

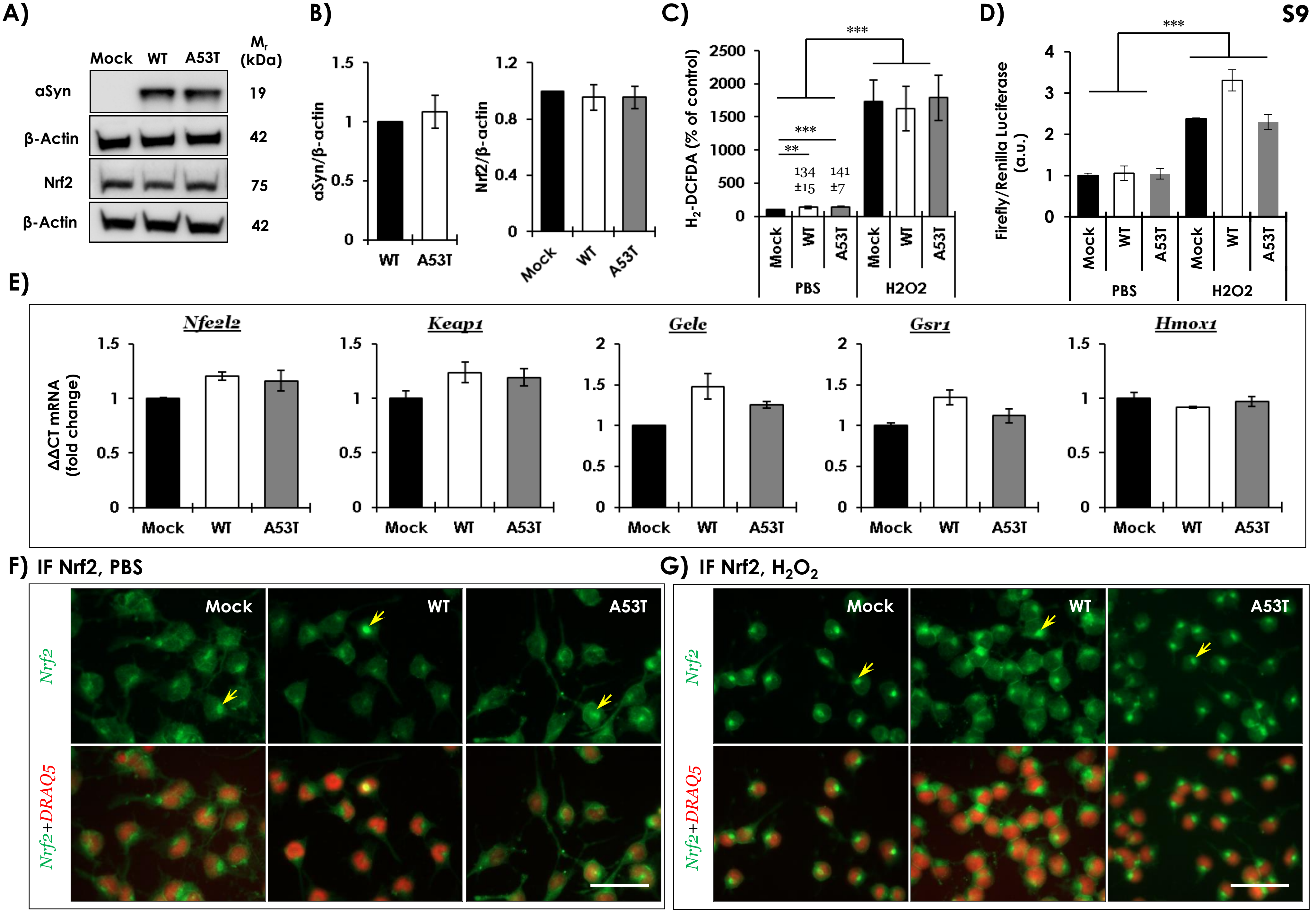

S9
A)
Mock WT A53T
Mr (kDa)
aSyn 19
β-Actin 42
Nrf2 75
β-Actin 42
B)
aSyn/β-actin
WT A53T
Nrf2/β-actin
Mock WT A53T
C)
H2-DCFDA (% of control)
***
***
**
134 ±15
141 ±7
Mock WT A53T Mock WT A53T
PBS H2O2
D)
Firefly/Renilla Luciferase (a.u.)
***
Mock WT A53T Mock WT A53T
PBS H2O2
E)
ΔΔCT mRNA (fold change)
Nfe2l2
Keap1
Gclc
Gsr1
Hmox1
Mock WT A53T
F) IF Nrf2, PBS
G) IF Nrf2, H2O2
Nrf2
Nrf2+DRAQ5
Mock WT A53T

**Table S1. Demographics, Control and PD cases.**

| **Case ID** | **Gender** | **Age (years)** | |
|---|---|---|---|
| **Controls** | | | |
| Control-1 | Male | 73 | |
| Control-2 | Male | 52 | |
| Control-3 | Female | 66 | |
| **PD** | | | **Additional postmortem pathology** |
| PD-1 | Male | 65 | Cortical Lewy body disease (LBD) |
| PD-2 | Male | 69 | Limbic predominant LBD |
| PD-3 | Male | 81 | Cortical LBD |
| PD-4 | Male | 74 | Rare lewy bodies in limbic and neocortex |
| PD-5 | Female | 68 | Cortical LBD |

| Table S2. Microarray datasets: Samples, genes and unique probe IDs | | | | |
|---|---|---|---|---|
| **GEO accession** | **GSE7621** | **GSE43490** | **GSE20146** | **GSE26927** |
| **Samples** | SN: Ctrl (n=9), PD (n=16) | SN: Ctrl (n=6), PD (n=8)<br>DMX: Ctrl (n=5), PD (n=8)<br>LC: Ctrl (n=7), PD (n=8) | GPi : Ctrl (n=10), PD (n=10) | SN : Ctrl (n=7), PD (n=12)<br>Ent. Ctx: Ctrl (n=6), AD (n=12)<br>Cerv. SC: Ctrl (n=7), ALS (n=9)<br>Caudate: Ctrl (n=10), HD (n=10)<br>Front. gyri: Ctrl (n=10), MS (n=10) |
| **Platform** | GPL570 Affymetrix Human Genome U133 | GPL6480 Agilent-014850 Whole Human Genome | GPL570 Affymetrix Human Genome U133 Plus | GPL6255 Illumina humanRef-8 v2.0 |
| **Reference** | *Lesnick, T.G. et al.,* | *Corradini, B.R. et al.,* | *Zheng, B. et al.,* | *Durrenberger, P.F. et al.,* |
| **Gene symbol** | **Unique Probe IDs** | | | |
| *NFE2L2* | 1567013_at | A_23_P5761 | 1567013_at | ILMN_9669 |
| *KEAP1* | 202417_at | A_23_P119141 | 202417_at | ILMN_18799 |
| *GSR* | 205770_at | A_23_P146084 | 205770_at | ILMN_14467 |
| *GCLC* | 1555330_at | A_23_P145114 | 1555330_at | ILMN_10857 |
| *HMOX1* | 203665_at | A_23_P120883 | 203665_at | ILMN_25059 |
| *TXN* | 208864_s_at | A_23_P60248 | 208864_s_at | ILMN_16294 |
| *CASP6* | 211464_x_at | A_23_P500799 | 211464_x_at | ILMN_11438 |
| *TNF* | 207113_s_at | Not found | 207113_s_at | Not found |
| *IL1A* | 208200_at | Not found | 208200_at | ILMN_25320 |
| *CASP3* | 202763_at | A_23_P92410 | 202763_at | ILMN_29066 |
| *NQO1* | 210519_s_at | Not found | 210519_s_at | ILMN_27575 |
| *PARK7* | 200006_at | A_23_P74740 | 200006_at | ILMN_6178 |

*Abbreviations:* **GSE7621**- Ctrl (control), PD (Parkinson Disease), SN (*substantia nigra*). **GSE43490-** Ctrl (control), PD (Parkinson Disease), SN (*substantia nigra*), DMX (*dorsal motor nucleus CN. X*), LC (*locus coeruleus*). **GSE20146-** Ctrl (control), PD (Parkinson Disease), GPi (*globus pallidus, interna*). **GSE26927-** Ctrl (control), PD (Parkinson Disease), AD (Alzheimer Disease), HD (Huntington Disease), ALS (Amyotrophic Lateral Sclerosis), MS (Multiple Sclerosis), SZ (Schizophrenia), SN (*substantia nigra*), Ent. Ctx (*entorhinal cortex*), Cerv. SC (*cervical spinal cord*), Front. Gyri (grey matter lesions in *frontal gyri*). Source: https://www.ncbi.nlm.nih.gov/geo/

| Table S3. Primer pairs used in qRT-PCR (mouse tissue/cells) | | |
|---|---|---|
| **Gene symbol** | **Forward** | **Reverse** |
| *Nfe2l2* | CACATTCCCAAACAAGATGCCT | TATCCAGGGCAAGCGACTCA |
| *Keap1* | GATCTACGTCCTCGGAGGCT | TCACTGTCCGGGTCATAGCA |
| *Hmox1* | TGCTAGCCTGGTGCAAGATAC | TGTCTGGGATGAGCTAGTGC |
| *Txn* | ATGACTGCCAGGATGTTGCT | AGAACTCCCCCACCTTTTGAC |
| *Gsr1* | GGGGCTCACTGAAGACGAAG | TCACAGCGTGATACATCGGG |
| *Gclc* | ACATCTACCACGCAGTCAAGG | CCAACATGTACTCCACCTCG |
| *Nqo1* | GGTAGCGGCTCCATGTACTC | CGCAGGATGCCACTCTGAAT |
| *Park7* | GCGGCTGCAGTCTTTAAGAAA | CAGTGACTTTGATCCCGGCT |
| *Tnf* | GTAGCCCACGTCGTAGCAAA | TTGAGATCCATGCCGTTGGC |
| *Il1* | CGCTTGAGTCGGCAAAGAAAT | TGGCAGAACTGTAGTCTTCGT |
| *Casp6* | AACCTGACTCGCAGGTTTTCA | GTTCTTCTGCTCTGAGGTCGT |
| *Casp3* | AGCTTGGAACGGTACGCTAA | GAGTCCACTGACTTGCTCCC |
| *Gapdh* | CCCTTAAGAGGGATGCTGCC | TACGGCCAAATCCGTTCACA |
